## Supplemental Data 1 for "Soil depth determines the microbial communities in *Sorghum bicolor* fields"

| sample_name | sex | race | refs | field | title | empo_1 | empo_2 | empo_3 | tube_id | altitude | depth_cm | genotype | [illegible] |
|---|---|---|---|---|---|---|---|---|---|---|---|---|---|
| 14937.9000I0001 | not applicable | Caudatum (Feterita) | [illegible] | D5 | sorghum soil with danforth | Free-living | Non-saline | Soil (non-saline) | NaN | not applicable | 15 | PI 656015 | [illegible] |
| 14937.9000I0002 | not applicable | Caudatum (Feterita) | [illegible] | D5 | sorghum soil with danforth | Free-living | Non-saline | Soil (non-saline) | NaN | not applicable | 30 | PI 656015 | [illegible] |
| 14937.9000I0003 | not applicable | Caudatum (Feterita) | [illegible] | D5 | sorghum soil with danforth | Free-living | Non-saline | Soil (non-saline) | NaN | not applicable | 60 | PI 656015 | [illegible] |
| 14937.9000I0004 | not applicable | Caudatum (Feterita) | [illegible] | D5 | sorghum soil with danforth | Free-living | Non-saline | Soil (non-saline) | NaN | not applicable | 90 | PI 656015 | [illegible] |
| 14937.9000I0005 | not applicable | Caudatum (Feterita) | [illegible] | D5 | sorghum soil with danforth | Free-living | Non-saline | Soil (non-saline) | NaN | not applicable | 120 | PI 656015 | [illegible] |
| 14937.9000I0006 | not applicable | Caudatum (Feterita) | [illegible] | D5 | sorghum soil with danforth | Free-living | Non-saline | Soil (non-saline) | NaN | not applicable | 150 | PI 656015 | [illegible] |
| 14937.9000I0007 | not applicable | Caudatum | [illegible] | D5 | sorghum soil with danforth | Free-living | Non-saline | Soil (non-saline) | NaN | not applicable | 15 | PI 595780 | [illegible] |
| 14937.9000I0008 | not applicable | Caudatum | [illegible] | D5 | sorghum soil with danforth | Free-living | Non-saline | Soil (non-saline) | NaN | not applicable | 30 | PI 595780 | [illegible] |
| 14937.9000I0009 | not applicable | Caudatum | [illegible] | D5 | sorghum soil with danforth | Free-living | Non-saline | Soil (non-saline) | NaN | not applicable | 60 | PI 595780 | [illegible] |
| 14937.9000I0010 | not applicable | Caudatum | [illegible] | D5 | sorghum soil with danforth | Free-living | Non-saline | Soil (non-saline) | NaN | not applicable | 90 | PI 595780 | [illegible] |
| 14937.9000I0011 | not applicable | Caudatum | [illegible] | D5 | sorghum soil with danforth | Free-living | Non-saline | Soil (non-saline) | NaN | not applicable | 120 | PI 595780 | [illegible] |
| 14937.9000I0012 | not applicable | Caudatum | [illegible] | D5 | sorghum soil with danforth | Free-living | Non-saline | Soil (non-saline) | NaN | not applicable | 150 | PI 595780 | [illegible] |
| 14937.9000I0013 | not applicable | Guinea | [illegible] | D5 | sorghum soil with danforth | Free-living | Non-saline | Soil (non-saline) | NaN | not applicable | 15 | PI 576434 | [illegible] |
| 14937.9000I0014 | not applicable | Guinea | [illegible] | D5 | sorghum soil with danforth | Free-living | Non-saline | Soil (non-saline) | NaN | not applicable | 30 | PI 576434 | [illegible] |
| 14937.9000I0015 | not applicable | Guinea | [illegible] | D5 | sorghum soil with danforth | Free-living | Non-saline | Soil (non-saline) | NaN | not applicable | 60 | PI 576434 | [illegible] |
| 14937.9000I0016 | not applicable | Guinea | [illegible] | D5 | sorghum soil with danforth | Free-living | Non-saline | Soil (non-saline) | NaN | not applicable | 90 | PI 576434 | [illegible] |
| 14937.9000I0017 | not applicable | Guinea | [illegible] | D5 | sorghum soil with danforth | Free-living | Non-saline | Soil (non-saline) | NaN | not applicable | 120 | PI 576434 | [illegible] |
| 14937.9000I0018 | not applicable | Guinea | [illegible] | D5 | sorghum soil with danforth | Free-living | Non-saline | Soil (non-saline) | NaN | not applicable | 150 | PI 576434 | [illegible] |
| 14937.9000I0019 | not applicable | Caudatum-Bicolor | [illegible] | D5 | sorghum soil with danforth | Free-living | Non-saline | Soil (non-saline) | NaN | not applicable | 15 | PI 597080 | [illegible] |
| 14937.9000I0020 | not applicable | Caudatum-Bicolor | [illegible] | D5 | sorghum soil with danforth | Free-living | Non-saline | Soil (non-saline) | NaN | not applicable | 30 | PI 597080 | [illegible] |
| 14937.9000I0021 | not applicable | Caudatum-Bicolor | [illegible] | D5 | sorghum soil with danforth | Free-living | Non-saline | Soil (non-saline) | NaN | not applicable | 60 | PI 597080 | [illegible] |
| 14937.9000I0022 | not applicable | Caudatum-Bicolor | [illegible] | D5 | sorghum soil with danforth | Free-living | Non-saline | Soil (non-saline) | NaN | not applicable | 90 | PI 597080 | [illegible] |
| 14937.9000I0023 | not applicable | Caudatum-Bicolor | [illegible] | D5 | sorghum soil with danforth | Free-living | Non-saline | Soil (non-saline) | NaN | not applicable | 120 | PI 597080 | [illegible] |
| 14937.9000I0024 | not applicable | Caudatum-Bicolor | [illegible] | D5 | sorghum soil with danforth | Free-living | Non-saline | Soil (non-saline) | NaN | not applicable | 150 | PI 597080 | [illegible] |
| 14937.9000I0025 | not applicable | Caudatum | [illegible] | D5 | sorghum soil with danforth | Free-living | Non-saline | Soil (non-saline) | NaN | not applicable | 15 | PI 656024 | [illegible] |
| 14937.9000I0026 | not applicable | Caudatum | [illegible] | D5 | sorghum soil with danforth | Free-living | Non-saline | Soil (non-saline) | NaN | not applicable | 30 | PI 656024 | [illegible] |
| 14937.9000I0027 | not applicable | Caudatum | [illegible] | D5 | sorghum soil with danforth | Free-living | Non-saline | Soil (non-saline) | NaN | not applicable | 60 | PI 656024 | [illegible] |
| 14937.9000I0028 | not applicable | Caudatum | [illegible] | D5 | sorghum soil with danforth | Free-living | Non-saline | Soil (non-saline) | NaN | not applicable | 90 | PI 656024 | [illegible] |
| 14937.9000I0029 | not applicable | Caudatum | [illegible] | D5 | sorghum soil with danforth | Free-living | Non-saline | Soil (non-saline) | NaN | not applicable | 120 | PI 656024 | [illegible] |
| 14937.9000I0030 | not applicable | Caudatum | [illegible] | D5 | sorghum soil with danforth | Free-living | Non-saline | Soil (non-saline) | NaN | not applicable | 150 | PI 656024 | [illegible] |
| 14937.9000I0031 | not applicable | Guinea-caudatum | [illegible] | D5 | sorghum soil with danforth | Free-living | Non-saline | Soil (non-saline) | NaN | not applicable | 15 | PI 595743 | [illegible] |
| 14937.9000I0032 | not applicable | Guinea-caudatum | [illegible] | D5 | sorghum soil with danforth | Free-living | Non-saline | Soil (non-saline) | NaN | not applicable | 30 | PI 595743 | [illegible] |
| 14937.9000I0033 | not applicable | Guinea-caudatum | [illegible] | D5 | sorghum soil with danforth | Free-living | Non-saline | Soil (non-saline) | NaN | not applicable | 60 | PI 595743 | [illegible] |
| 14937.9000I0034 | not applicable | Guinea-caudatum | [illegible] | D5 | sorghum soil with danforth | Free-living | Non-saline | Soil (non-saline) | NaN | not applicable | 90 | PI 595743 | [illegible] |
| 14937.9000I0035 | not applicable | Guinea-caudatum | [illegible] | D5 | sorghum soil with danforth | Free-living | Non-saline | Soil (non-saline) | NaN | not applicable | 120 | PI 595743 | [illegible] |
| 14937.9000I0036 | not applicable | Guinea-caudatum | [illegible] | D5 | sorghum soil with danforth | Free-living | Non-saline | Soil (non-saline) | NaN | not applicable | 150 | PI 595743 | [illegible] |
| 14937.9000I0037 | not applicable | Guinea (Conspicuum) | [illegible] | D5 | sorghum soil with danforth | Free-living | Non-saline | Soil (non-saline) | NaN | not applicable | 15 | PI 533766 | [illegible] |
| 14937.9000I0038 | not applicable | Guinea (Conspicuum) | [illegible] | D5 | sorghum soil with danforth | Free-living | Non-saline | Soil (non-saline) | NaN | not applicable | 30 | PI 533766 | [illegible] |
| [illegible] | [illegible] | [illegible] | [illegible] | [illegible] | [illegible] | [illegible] | [illegible] | [illegible] | [illegible] | [illegible] | [illegible] | [illegible] | [illegible] |

| | | | | | | | | | | | | | | | | | | | | | | | | | | | | | | | | | | | | | | | | | | | | | | | | | |
|---|---|---|---|---|---|---|---|---|---|---|---|---|---|---|---|---|---|---|---|---|---|---|---|---|---|---|---|---|---|---|---|---|---|---|---|---|---|---|---|---|---|---|---|---|---|---|---|---|---|
| 14937.5000E0139 | not applicable | Caudatum-Durra | https://academic.oup.com/plphys/article/182/2/977/6116230[illegible] | DS | sorghum soil with [illegible] | Free-living | Non-saline | Soil (non-saline) | NaN | not applicable | 60 | PI 533762 | not applicable | [illegible] | [illegible] | 135 | urban biome | -90.68664 | largest root area (2020 RSA study) | 37.7 | Tamboro 7 | Sudan | not applicable | TBD | farmland soil | soil | not applicable | not applicable | | 2 | 0.279 | DS 1193_1194 TOP DEPTHS | soil | 5.7 | soil | USA:MO: StLouis | 0 | 24 | 23 | | TRUE | not applicable | not applicable | 24.06 | DS 1193/1194 60 | not applicable | DS__1193_1194 | soil metagenome | [illegible] |
| 14937.5000E0140 | not applicable | Caudatum-Durra | https://academic.oup.com/plphys/article/182/2/977/6116230[illegible] | DS | sorghum soil with [illegible] | Free-living | Non-saline | Soil (non-saline) | NaN | not applicable | 90 | PI 533762 | not applicable | [illegible] | [illegible] | 135 | urban biome | -90.68664 | largest root area (2020 RSA study) | 13.2 | Tamboro 7 | Sudan | not applicable | TBD | farmland soil | soil | not applicable | not applicable | | 2 | 0.204 | DS 1193_1194 LOWER DEPTHS | soil | 4.8 | soil | USA:MO: StLouis | 24 | 60 | 22 | | TRUE | not applicable | not applicable | 24 | DS 1193/1194 90 | not applicable | DS__1193_1194 | soil metagenome | [illegible] |
| 14937.5000E0141 | not applicable | Caudatum-Durra | https://academic.oup.com/plphys/article/182/2/977/6116230[illegible] | DS | sorghum soil with [illegible] | Free-living | Non-saline | Soil (non-saline) | NaN | not applicable | 120 | PI 533762 | not applicable | [illegible] | [illegible] | 135 | urban biome | -90.68664 | largest root area (2020 RSA study) | 13.2 | Tamboro 7 | Sudan | not applicable | TBD | farmland soil | soil | not applicable | not applicable | | 2 | 0.204 | DS 1193_1194 LOWER DEPTHS | soil | 4.8 | soil | USA:MO: StLouis | 24 | 60 | 22 | | TRUE | not applicable | not applicable | 24 | DS 1193/1194 120 | not applicable | DS__1193_1194 | soil metagenome | [illegible] |
| 14937.5000E0142 | not applicable | Caudatum-Durra | https://academic.oup.com/plphys/article/182/2/977/6116230[illegible] | DS | sorghum soil with [illegible] | Free-living | Non-saline | Soil (non-saline) | NaN | not applicable | 150 | PI 533762 | not applicable | [illegible] | [illegible] | 135 | urban biome | -90.68664 | largest root area (2020 RSA study) | 13.2 | Tamboro 7 | Sudan | not applicable | TBD | farmland soil | soil | not applicable | not applicable | | 2 | 0.204 | DS 1193_1194 LOWER DEPTHS | soil | 4.8 | soil | USA:MO: StLouis | 24 | 60 | 22 | | TRUE | not applicable | not applicable | 24 | DS 1193/1194 150 | not applicable | DS__1193_1194 | soil metagenome | [illegible] |
| 14937.5000E0143 | not applicable | Guinea-caudatum | https://www.nature.com/articles/s41598-017-18119-1 | DS | sorghum soil with [illegible] | Free-living | Non-saline | Soil (non-saline) | NaN | not applicable | 15 | PI 595743 | not applicable | [illegible] | [illegible] | 135 | urban biome | -90.68664 | bmr phenotype, mutation in SbCAD2 | 34.1 | SAP-154 | United States | not applicable | TBD | farmland soil | soil | not applicable | not applicable | Insensitive | 2 | 0.92 | DS 1271_1272 TOP DEPTHS | soil | 5.6 | soil | USA:MO: StLouis | 0 | 24 | 25 | Forage | TRUE | not applicable | not applicable | 23.48 | DS 1271/1272 15 | not applicable | DS__1271_1272 | soil metagenome | [illegible] |
| 14937.5000E0144 | not applicable | Guinea-caudatum | https://www.nature.com/articles/s41598-017-18119-1 | DS | sorghum soil with [illegible] | Free-living | Non-saline | Soil (non-saline) | NaN | not applicable | 30 | PI 595743 | not applicable | [illegible] | [illegible] | 135 | urban biome | -90.68664 | bmr phenotype, mutation in SbCAD2 | 34.1 | SAP-154 | United States | not applicable | TBD | farmland soil | soil | not applicable | not applicable | Insensitive | 2 | 0.92 | DS 1271_1272 TOP DEPTHS | soil | 5.6 | soil | USA:MO: StLouis | 0 | 24 | 25 | Forage | TRUE | not applicable | not applicable | 23.48 | DS 1271/1272 30 | not applicable | DS__1271_1272 | soil metagenome | [illegible] |
| 14937.5000E0145 | not applicable | Guinea-caudatum | https://www.nature.com/articles/s41598-017-18119-1 | DS | sorghum soil with [illegible] | Free-living | Non-saline | Soil (non-saline) | NaN | not applicable | 60 | PI 595743 | not applicable | [illegible] | [illegible] | 135 | urban biome | -90.68664 | bmr phenotype, mutation in SbCAD2 | 34.1 | SAP-154 | United States | not applicable | TBD | farmland soil | soil | not applicable | not applicable | Insensitive | 2 | 0.92 | DS 1271_1272 TOP DEPTHS | soil | 5.6 | soil | USA:MO: StLouis | 0 | 24 | 25 | Forage | TRUE | not applicable | not applicable | 23.48 | DS 1271/1272 60 | not applicable | DS__1271_1272 | soil metagenome | [illegible] |
| 14937.5000E0146 | not applicable | Guinea-caudatum | https://www.nature.com/articles/s41598-017-18119-1 | DS | sorghum soil with [illegible] | Free-living | Non-saline | Soil (non-saline) | NaN | not applicable | 90 | PI 595743 | not applicable | [illegible] | [illegible] | 135 | urban biome | -90.68664 | bmr phenotype, mutation in SbCAD2 | 8.4 | SAP-154 | United States | not applicable | TBD | farmland soil | soil | not applicable | not applicable | Insensitive | 2 | 0.118 | DS 1271_1272 LOWER DEPTHS | soil | 4.8 | soil | USA:MO: StLouis | 24 | 60 | 28 | Forage | TRUE | not applicable | not applicable | 31.82 | DS 1271/1272 90 | not applicable | DS__1271_1272 | soil metagenome | [illegible] |
| 14937.5000E0147 | not applicable | Guinea-caudatum | https://www.nature.com/articles/s41598-017-18119-1 | DS | sorghum soil with [illegible] | Free-living | Non-saline | Soil (non-saline) | NaN | not applicable | 120 | PI 595743 | not applicable | [illegible] | [illegible] | 135 | urban biome | -90.68664 | bmr phenotype, mutation in SbCAD2 | 8.4 | SAP-154 | United States | not applicable | TBD | farmland soil | soil | not applicable | not applicable | Insensitive | 2 | 0.118 | DS 1271_1272 LOWER DEPTHS | soil | 4.8 | soil | USA:MO: StLouis | 24 | 60 | 28 | Forage | TRUE | not applicable | not applicable | 31.82 | DS 1271/1272 120 | not applicable | DS__1271_1272 | soil metagenome | [illegible] |
| 14937.5000E0148 | not applicable | Guinea-caudatum | https://www.nature.com/articles/s41598-017-18119-1 | DS | sorghum soil with [illegible] | Free-living | Non-saline | Soil (non-saline) | NaN | not applicable | 150 | PI 595743 | not applicable | [illegible] | [illegible] | 135 | urban biome | -90.68664 | bmr phenotype, mutation in SbCAD2 | 8.4 | SAP-154 | United States | not applicable | TBD | farmland soil | soil | not applicable | not applicable | Insensitive | 2 | 0.118 | DS 1271_1272 LOWER DEPTHS | soil | 4.8 | soil | USA:MO: StLouis | 24 | 60 | 28 | Forage | TRUE | not applicable | not applicable | 31.82 | DS 1271/1272 150 | not applicable | DS__1271_1272 | soil metagenome | [illegible] |
| 14937.5000E0149 | not applicable | Guinea (Conspicuum) | https://academic.oup.com/genetics/article/206/2/573/6064253?login=true | DS | sorghum soil with [illegible] | Free-living | Non-saline | Soil (non-saline) | NaN | not applicable | 15 | PI 533869 | not applicable | [illegible] | [illegible] | 135 | urban biome | -90.68664 | Grain NAM Parent | 31.9 | SC283 | Tanzania | not applicable | TBD | farmland soil | soil | not applicable | not applicable | | 1 | 1.075 | DS 1367_1368 TOP DEPTHS | soil | 5.5 | soil | USA:MO: StLouis | 0 | 24 | 17 | Grain | TRUE | not applicable | not applicable | 26.05 | DS 1367/1368 15 | not applicable | DS__1367_1368 | soil metagenome | [illegible] |
| 14937.5000E0150 | not applicable | Guinea (Conspicuum) | https://academic.oup.com/genetics/article/206/2/573/6064253?login=true | DS | sorghum soil with [illegible] | Free-living | Non-saline | Soil (non-saline) | NaN | not applicable | 30 | PI 533869 | not applicable | [illegible] | [illegible] | 135 | urban biome | -90.68664 | Grain NAM Parent | 31.9 | SC283 | Tanzania | not applicable | TBD | farmland soil | soil | not applicable | not applicable | | 1 | 1.075 | DS 1367_1368 TOP DEPTHS | soil | 5.5 | soil | USA:MO: StLouis | 0 | 24 | 17 | Grain | TRUE | not applicable | not applicable | 26.05 | DS 1367/1368 30 | not applicable | DS__1367_1368 | soil metagenome | [illegible] |
| 14937.5000E0151 | not applicable | Guinea (Conspicuum) | https://academic.oup.com/genetics/article/206/2/573/6064253?login=true | DS | sorghum soil with [illegible] | Free-living | Non-saline | Soil (non-saline) | NaN | not applicable | 60 | PI 533869 | not applicable | [illegible] | [illegible] | 135 | urban biome | -90.68664 | Grain NAM Parent | 31.9 | SC283 | Tanzania | not applicable | TBD | farmland soil | soil | not applicable | not applicable | | 1 | 1.075 | DS 1367_1368 TOP DEPTHS | soil | 5.5 | soil | USA:MO: StLouis | 0 | 24 | 17 | Grain | TRUE | not applicable | not applicable | 26.05 | DS 1367/1368 60 | not applicable | DS__1367_1368 | soil metagenome | [illegible] |
| 14937.5000E0152 | not applicable | Guinea (Conspicuum) | https://academic.oup.com/genetics/article/206/2/573/6064253?login=true | DS | sorghum soil with [illegible] | Free-living | Non-saline | Soil (non-saline) | NaN | not applicable | 90 | PI 533869 | not applicable | [illegible] | [illegible] | 135 | urban biome | -90.68664 | Grain NAM Parent | 11.1 | SC283 | Tanzania | not applicable | TBD | farmland soil | soil | not applicable | not applicable | | 1 | 0.033 | DS 1367_1368 LOWER DEPTHS | soil | 4.8 | soil | USA:MO: StLouis | 24 | 60 | 39 | Grain | TRUE | not applicable | not applicable | 32.86 | DS 1367/1368 90 | not applicable | DS__1367_1368 | soil metagenome | [illegible] |
| 14937.5000E0153 | not applicable | Guinea (Conspicuum) | https://academic.oup.com/genetics/article/206/2/573/6064253?login=true | DS | sorghum soil with [illegible] | Free-living | Non-saline | Soil (non-saline) | NaN | not applicable | 120 | PI 533869 | not applicable | [illegible] | [illegible] | 135 | urban biome | -90.68664 | Grain NAM Parent | 11.1 | SC283 | Tanzania | not applicable | TBD | farmland soil | soil | not applicable | not applicable | | 1 | 0.033 | DS 1367_1368 LOWER DEPTHS | soil | 4.8 | soil | USA:MO: StLouis | 24 | 60 | 39 | Grain | TRUE | not applicable | not applicable | 32.86 | DS 1367/1368 120 | not applicable | DS__1367_1368 | soil metagenome | [illegible] |
| 14937.5000E0154 | not applicable | Guinea (Conspicuum) | https://academic.oup.com/genetics/article/206/2/573/6064253?login=true | DS | sorghum soil with [illegible] | Free-living | Non-saline | Soil (non-saline) | NaN | not applicable | 150 | PI 533869 | not applicable | [illegible] | [illegible] | 135 | urban biome | -90.68664 | Grain NAM Parent | 11.1 | SC283 | Tanzania | not applicable | TBD | farmland soil | soil | not applicable | not applicable | | 1 | 0.033 | DS 1367_1368 LOWER DEPTHS | soil | 4.8 | soil | USA:MO: StLouis | 24 | 60 | 39 | Grain | TRUE | not applicable | not applicable | 32.86 | DS 1367/1368 150 | not applicable | DS__1367_1368 | soil metagenome | [illegible] |
| [illegible] | | | | | | | | | | | | | | | | | | | | | | | | | | | | | | | | | | | | | | | | | | | | | | | | | |

| | | | | | | | | | | | | | | | | | | | | | | | |
|---|---|---|---|---|---|---|---|---|---|---|---|---|---|---|---|---|---|---|---|---|---|---|---|
| 14937.[illegible] | not applicable | Durra | https://academic.oup.com/[illegible] | A3 | sorghum soil with [illegible] | Free-living | Non-saline | Soil (non-saline) | NaN | not applicable | 30 | PI 329311 | not applicable | [illegible] | [illegible] | 135 | urban biome | -90.68742 | Carbon Partitioning NAM Parent | 45.1 | IS 11069 | Ethiopia | [illegible] |
| 14937.[illegible] | not applicable | Durra | https://academic.oup.com/[illegible] | A3 | sorghum soil with [illegible] | Free-living | Non-saline | Soil (non-saline) | NaN | not applicable | 60 | PI 329311 | not applicable | [illegible] | [illegible] | 135 | urban biome | -90.68742 | Carbon Partitioning NAM Parent | 45.1 | IS 11069 | Ethiopia | [illegible] |
| 14937.[illegible] | not applicable | Durra | https://academic.oup.com/[illegible] | A3 | sorghum soil with [illegible] | Free-living | Non-saline | Soil (non-saline) | NaN | not applicable | 90 | PI 329311 | not applicable | [illegible] | [illegible] | 135 | urban biome | -90.68742 | Carbon Partitioning NAM Parent | 13.9 | IS 11069 | Ethiopia | [illegible] |
| 14937.[illegible] | not applicable | Durra | https://academic.oup.com/[illegible] | A3 | sorghum soil with [illegible] | Free-living | Non-saline | Soil (non-saline) | NaN | not applicable | 120 | PI 329311 | not applicable | [illegible] | [illegible] | 135 | urban biome | -90.68742 | Carbon Partitioning NAM Parent | 13.9 | IS 11069 | Ethiopia | [illegible] |
| 14937.[illegible] | not applicable | Durra | https://academic.oup.com/[illegible] | A3 | sorghum soil with [illegible] | Free-living | Non-saline | Soil (non-saline) | NaN | not applicable | 150 | PI 329311 | not applicable | [illegible] | [illegible] | 135 | urban biome | -90.68742 | Carbon Partitioning NAM Parent | 13.9 | IS 11069 | Ethiopia | [illegible] |
| 14937.[illegible] | not applicable | Bicolor | https://academic.oup.com/[illegible] | A3 | sorghum soil with [illegible] | Free-living | Non-saline | Soil (non-saline) | NaN | not applicable | 15 | PI22913 | not applicable | [illegible] | [illegible] | 135 | urban biome | -90.68742 | Carbon Partitioning NAM Parent | 77.9 | Chinese Amber | China | [illegible] |
| 14937.[illegible] | not applicable | Bicolor | https://academic.oup.com/[illegible] | A3 | sorghum soil with [illegible] | Free-living | Non-saline | Soil (non-saline) | NaN | not applicable | 30 | PI22913 | not applicable | [illegible] | [illegible] | 135 | urban biome | -90.68742 | Carbon Partitioning NAM Parent | 77.9 | Chinese Amber | China | [illegible] |
| 14937.[illegible] | not applicable | Bicolor | https://academic.oup.com/[illegible] | A3 | sorghum soil with [illegible] | Free-living | Non-saline | Soil (non-saline) | NaN | not applicable | 60 | PI22913 | not applicable | [illegible] | [illegible] | 135 | urban biome | -90.68742 | Carbon Partitioning NAM Parent | 77.9 | Chinese Amber | China | [illegible] |
| 14937.[illegible] | not applicable | Bicolor | https://academic.oup.com/[illegible] | A3 | sorghum soil with [illegible] | Free-living | Non-saline | Soil (non-saline) | NaN | not applicable | 90 | PI22913 | not applicable | [illegible] | [illegible] | 135 | urban biome | -90.68742 | Carbon Partitioning NAM Parent | 17.3 | Chinese Amber | China | [illegible] |
| 14937.[illegible] | not applicable | Bicolor | https://academic.oup.com/[illegible] | A3 | sorghum soil with [illegible] | Free-living | Non-saline | Soil (non-saline) | NaN | not applicable | 120 | PI22913 | not applicable | [illegible] | [illegible] | 135 | urban biome | -90.68742 | Carbon Partitioning NAM Parent | 17.3 | Chinese Amber | China | [illegible] |
| 14937.[illegible] | not applicable | Bicolor | https://academic.oup.com/[illegible] | A3 | sorghum soil with [illegible] | Free-living | Non-saline | Soil (non-saline) | NaN | not applicable | 150 | PI22913 | not applicable | [illegible] | [illegible] | 135 | urban biome | -90.68742 | Carbon Partitioning NAM Parent | 17.3 | Chinese Amber | China | [illegible] |
| 14937.[illegible] | not applicable | Caudatum | https://www.sciencedirect.com/[illegible] | A3 | sorghum soil with [illegible] | Free-living | Non-saline | Soil (non-saline) | NaN | not applicable | 15 | PI 154844 | not applicable | [illegible] | [illegible] | 135 | urban biome | -90.68742 | Carbon Partitioning NAM Parent | 53.9 | Grassl | Uganda | [illegible] |
| 14937.[illegible] | not applicable | Caudatum | https://www.sciencedirect.com/[illegible] | A3 | sorghum soil with [illegible] | Free-living | Non-saline | Soil (non-saline) | NaN | not applicable | 30 | PI 154844 | not applicable | [illegible] | [illegible] | 135 | urban biome | -90.68742 | Carbon Partitioning NAM Parent | 53.9 | Grassl | Uganda | [illegible] |
| 14937.[illegible] | not applicable | Caudatum | https://www.sciencedirect.com/[illegible] | A3 | sorghum soil with [illegible] | Free-living | Non-saline | Soil (non-saline) | NaN | not applicable | 60 | PI 154844 | not applicable | [illegible] | [illegible] | 135 | urban biome | -90.68742 | Carbon Partitioning NAM Parent | 53.9 | Grassl | Uganda | [illegible] |
| [illegible] | [illegible] | [illegible] | [illegible] | [illegible] | [illegible] | [illegible] | [illegible] | [illegible] | [illegible] | [illegible] | [illegible] | [illegible] | [illegible] | [illegible] | [illegible] | [illegible] | [illegible] | [illegible] | [illegible] | [illegible] | [illegible] | [illegible] | [illegible] |
