## Supplemental Data 2 for "Soil depth determines the microbial communities in *Sorghum bicolor* fields"

| | faith_pd | | shannon_entropy | | chao1 | |
|---|---|---|---|---|---|---|
| **Soil Property** | **Spearman Rho** | **P-Value** | **Spearman Rho.1** | **P-Value.1** | **Spearman Rho.2** | **P-Value.2** |
| depth_cm | -0.74 | <0.001*** | -0.784 | <0.001*** | -0.778 | <0.001*** |
| root_mass_g | 0.552 | <0.001*** | 0.545 | <0.001*** | 0.572 | <0.001*** |
| soi_h2o_total_organic_c | 0.523 | <0.001*** | 0.534 | <0.001*** | 0.523 | <0.001*** |
| soil_available_k | -0.586 | <0.001*** | -0.602 | <0.001*** | -0.609 | <0.001*** |
| soil_available_n | -0.392 | <0.001*** | -0.356 | <0.001*** | -0.401 | <0.001*** |
| soil_available_p | 0.25 | <0.001*** | 0.17 | <0.001*** | 0.204 | <0.001*** |
| soil_calcium_ppm_ca | -0.423 | <0.001*** | -0.475 | <0.001*** | -0.467 | <0.001*** |
| soil_co2c | 0.639 | <0.001*** | 0.616 | <0.001*** | 0.621 | <0.001*** |
| soil_composition_order | -0.734 | <0.001*** | -0.717 | <0.001*** | -0.737 | <0.001*** |
| soil_copper_ppm_cu | - | 0.02 | - | 0.015 | 0.169 | <0.001*** |
| soil_h2o_organic_n | 0.583 | <0.001*** | 0.569 | <0.001*** | 0.572 | <0.001*** |
| soil_h2o_total_n | 0.465 | <0.001*** | 0.47 | <0.001*** | 0.472 | <0.001*** |
| soil_h3a_ammonium | -0.446 | <0.001*** | -0.413 | <0.001*** | -0.446 | <0.001*** |
| soil_h3a_icap_aluminum | -0.635 | <0.001*** | -0.634 | <0.001*** | -0.635 | <0.001*** |
| soil_h3a_icap_calcium | 0.693 | <0.001*** | 0.636 | <0.001*** | 0.664 | <0.001*** |
| soil_h3a_icap_copper | -0.498 | <0.001*** | -0.452 | <0.001*** | -0.449 | <0.001*** |
| soil_h3a_icap_iron | -0.318 | <0.001*** | -0.362 | <0.001*** | -0.304 | <0.001*** |
| soil_h3a_icap_magnesium | -0.317 | <0.001*** | -0.372 | <0.001*** | -0.352 | <0.001*** |
| soil_h3a_icap_manganese | 0.688 | <0.001*** | 0.642 | <0.001*** | 0.65 | <0.001*** |
| soil_h3a_icap_potassium | -0.587 | <0.001*** | -0.602 | <0.001*** | -0.609 | <0.001*** |
| soil_h3a_icap_sodium | -0.648 | <0.001*** | -0.636 | <0.001*** | -0.635 | <0.001*** |
| soil_h3a_icap_sulfur | -0.476 | <0.001*** | -0.515 | <0.001*** | -0.516 | <0.001*** |
| soil_h3a_icap_zinc | 0.389 | <0.001*** | 0.307 | <0.001*** | 0.342 | <0.001*** |
| soil_h3a_inorganic_nitrogen | -0.195 | <0.001*** | - | 0.002 | -0.185 | <0.001*** |
| soil_h3a_inorganic_phosphorus | 0.213 | <0.001*** | - | 0.01 | - | 0.001 |
| soil_h3a_nitrate | 0.203 | <0.001*** | 0.225 | <0.001*** | 0.212 | <0.001*** |
| soil_h3a_organic_phosphorus | 0.412 | <0.001*** | 0.346 | <0.001*** | 0.375 | <0.001*** |
| soil_h3a_total_phosphorus | 0.245 | <0.001*** | 0.164 | 0.001 | 0.2 | <0.001*** |
| soil_haney_test_n | -0.392 | <0.001*** | -0.356 | <0.001*** | -0.401 | <0.001*** |
| soil_iron_ppm_fe | - | 0.742 | - | 0.59 | - | 0.134 |
| soil_lbs_n_a | - | 0.603 | - | 0.911 | - | 0.782 |
| soil_lbs_n_difference | -0.451 | <0.001*** | -0.423 | <0.001*** | -0.467 | <0.001*** |
| soil_magnesium_ppm_mg | -0.609 | <0.001*** | -0.632 | <0.001*** | -0.635 | <0.001*** |
| soil_manganese_ppm_mn | -0.42 | <0.001*** | -0.382 | <0.001*** | -0.416 | <0.001*** |
| soil_n_savings | -0.451 | <0.001*** | -0.423 | <0.001*** | -0.468 | <0.001*** |
| soil_nitrate_n_ppm_n | 0.234 | <0.001*** | 0.254 | <0.001*** | 0.244 | <0.001*** |
| soil_nutrient_value | - | 0.002 | -0.21 | <0.001*** | -0.203 | <0.001*** |
| soil_olsen_p_ppm_p | - | 0.104 | - | 0.874 | - | 0.485 |
| soil_organic_c_to_n_ratio | -0.384 | <0.001*** | -0.354 | <0.001*** | -0.364 | <0.001*** |
| soil_organic_n_release | 0.592 | <0.001*** | 0.578 | <0.001*** | 0.579 | <0.001*** |
| soil_organic_n_reserve | -0.264 | <0.001*** | -0.259 | <0.001*** | -0.25 | <0.001*** |
| soil_organic_n_to_inorganicn_ratio | 0.581 | <0.001*** | 0.554 | <0.001*** | 0.566 | <0.001*** |
| soil_organicmatter | 0.557 | <0.001*** | 0.513 | <0.001*** | 0.549 | <0.001*** |
| soil_percent_mac | 0.618 | <0.001*** | 0.583 | <0.001*** | 0.6 | <0.001*** |
| soil_ph_1v1 | 0.69 | <0.001*** | 0.635 | <0.001*** | 0.65 | <0.001*** |
| soil_potassium_ppm_k | -0.628 | <0.001*** | -0.608 | <0.001*** | -0.634 | <0.001*** |

| | | | | | | |
|---|---|---|---|---|---|---|
| soil_sodium_ppm_na | -0.676 | <0.001*** | -0.64 | <0.001*** | -0.65 | <0.001*** |
| soil_soil_health_calculation | 0.636 | <0.001*** | 0.616 | <0.001*** | 0.623 | <0.001*** |
| soil_soluble_salt_1v1 | -0.36 | <0.001*** | -0.316 | <0.001*** | -0.387 | <0.001*** |
| soil_sulfate_s_ppm_s | -0.598 | <0.001*** | -0.585 | <0.001*** | -0.614 | <0.001*** |
| soil_traditional_n | - | 0.784 | - | 0.419 | - | 0.665 |
| soil_wdrfbuffer | 0.682 | <0.001*** | 0.646 | <0.001*** | 0.651 | <0.001*** |
| soil_zinc_ppm_zn | 0.456 | <0.001*** | 0.422 | <0.001*** | 0.438 | <0.001*** |
