## Supplemental Data 3 for "Soil depth determines the microbial communities in *Sorghum bicolor* fields"

| Soil Property | Kruskal-Wallis | p-value |
|---|---|---|
| root_mass_g | 228.3557868 | 1.36E-51 |
| soi_h2o_total_organic_c | 206.5677142 | 7.70E-47 |
| soil_available_k | 260.4847042 | 1.35E-58 |
| soil_available_n | 97.1378575 | 6.47E-23 |
| soil_available_p | 15.07076529 | 1.04E-04 |
| soil_calcium_ppm_ca | 164.5286384 | 1.16E-37 |
| soil_co2c | 258.8204395 | 3.10E-58 |
| soil_composition_order | 380.809317 | 8.29E-85 |
| soil_copper_ppm_cu | 22.28289411 | 2.35E-06 |
| soil_h2o_organic_n | 222.5072055 | 2.57E-50 |
| soil_h2o_total_n | 160.6786184 | 8.04E-37 |
| soil_h3a_ammonium | 98.66259461 | 2.99E-23 |
| soil_h3a_icap_aluminum | 261.9531375 | 6.44E-59 |
| soil_h3a_icap_calcium | 260.0942679 | 1.64E-58 |
| soil_h3a_icap_copper | 128.0434808 | 1.10E-29 |
| soil_h3a_icap_iron | 96.62159728 | 8.39E-23 |
| soil_h3a_icap_magnesium | 109.8410947 | 1.06E-25 |
| soil_h3a_icap_manganese | 259.5005304 | 2.20E-58 |
| soil_h3a_icap_potassium | 260.9744327 | 1.05E-58 |
| soil_h3a_icap_sodium | 264.7055565 | 1.62E-59 |
| soil_h3a_icap_sulfur | 216.1473764 | 6.26E-49 |
| soil_h3a_icap_zinc | 54.76999318 | 1.35E-13 |
| soil_h3a_inorganic_nitrogen | 12.01118333 | 5.29E-04 |
| soil_h3a_inorganic_phosphorus | 9.701031506 | 1.84E-03 |
| soil_h3a_nitrate | 35.17858987 | 3.01E-09 |
| soil_h3a_organic_phosphorus | 57.53012677 | 3.33E-14 |
| soil_h3a_total_phosphorus | 14.18821506 | 1.65E-04 |
| soil_haney_test_n | 97.1378575 | 6.47E-23 |
| soil_iron_ppm_fe | 1.941432338 | 1.64E-01 |
| soil_lbs_n_a | 0.00796749 | 9.29E-01 |
| soil_lbs_n_difference | 133.5205624 | 6.96E-31 |
| soil_magnesium_ppm_mg | 270.3160168 | 9.68E-61 |
| soil_manganese_ppm_mn | 100.5501554 | 1.15E-23 |
| soil_n_savings | 133.2852709 | 7.83E-31 |
| soil_nitrate_n_ppm_n | 47.89652685 | 4.49E-12 |
| soil_nutrient_value | 39.07166231 | 4.09E-10 |
| soil_olsen_p_ppm_p | 0.09414762 | 7.59E-01 |
| soil_organic_c_to_n_ratio | 80.11901304 | 3.53E-19 |

| | | |
|---|---|---|
| soil_organic_n_release | 222.4359784 | 2.66E-50 |
| soil_organic_n_reserve | 30.53760237 | 3.27E-08 |
| soil_organic_n_to_inorganicn_rati | 200.2051295 | 1.88E-45 |
| soil_organicmatter | 183.8904097 | 6.86E-42 |
| soil_percent_mac | 236.6099017 | 2.16E-53 |
| soil_ph_1v1 | 252.8707514 | 6.15E-57 |
| soil_potassium_ppm_k | 241.2845774 | 2.06E-54 |
| soil_sodium_ppm_na | 261.140174 | 9.68E-59 |
| soil_soil_health_calculation | 263.5562079 | 2.88E-59 |
| soil_soluble_salt_1v1 | 90.43715255 | 1.91E-21 |
| soil_sulfate_s_ppm_s | 247.7813563 | 7.91E-56 |
| soil_traditional_n | 0.3702870506 | 5.43E-01 |
| soil_wdrfbuffer | 262.9797856 | 3.85E-59 |
| soil_zinc_ppm_zn | 132.6097252 | 1.10E-30 |
