## Supplemental Data 4 for "Soil depth determines the microbial communities in *Sorghum bicolor* fields"

| Feature ID | 2% | 9% | 25% | 50% | 75% | 91% | 98% |
|---|---|---|---|---|---|---|---|
| k__Bacteria;p__Chloroflexi;c__Ktedonobacteria;o__Thermogemmatisporales;f__Thermogemmatisporaceae | 0 | 0 | 1 | 76 | 224 | 347.96 | 511.48 |
| k__Bacteria;p__Acidobacteria;c__Acidobacteriia;o__Acidobacteriales;f__Koribacteraceae | 0 | 7 | 19 | 51 | 89 | 129.92 | 162.76 |
| k__Bacteria;p__Acidobacteria;c__Acidobacteria-6;o__iii1-15;f__ | 0.12 | 3 | 14 | 40 | 69 | 97 | 118.76 |
| k__Bacteria;p__Actinobacteria;c__Thermoleophilia;o__Gaiellales;f__Gaiellaceae | 6.24 | 17 | 26 | 37 | 52 | 71 | 100.88 |
| k__Bacteria;p__Verrucomicrobia;c__[Spartobacteria];o__[Chthoniobacterales];f__[Chthoniobacteraceae] | 0 | 4 | 17 | 33 | 54 | 87 | 132.76 |
| k__Bacteria;p__Proteobacteria;c__Alphaproteobacteria;o__Rhodospirillales;f__Rhodospirillaceae | 7 | 12 | 19 | 27 | 39 | 49.96 | 64.88 |
| k__Archaea;p__Crenarchaeota;c__Thaumarchaeota;o__Nitrososphaerales;f__Nitrososphaeraceae | 2 | 5 | 12 | 27 | 48 | 75.96 | 125.28 |
| k__Bacteria;p__AD3;c__ABS-6;o__;f__ | 0 | 0 | 0 | 25 | 117 | 225.92 | 360.76 |
| k__Bacteria;p__Proteobacteria;c__Deltaproteobacteria;o__Syntrophobacterales;f__Syntrophobacteraceae | 0 | 6 | 15 | 25 | 36 | 50 | 58.88 |
| k__Bacteria;p__Proteobacteria;c__Alphaproteobacteria;o__Rhizobiales;f__Hyphomicrobiaceae | 1 | 6 | 14 | 24 | 36 | 50 | 62 |
| k__Bacteria;p__Acidobacteria;c__[Chloracidobacteria];o__RB41;f__ | 0 | 1 | 6 | 20 | 39 | 62.96 | 90.88 |
| k__Bacteria;p__Gemmatimonadetes;c__Gemm-1;o__;f__ | 1 | 4 | 8 | 15 | 28 | 47.96 | 64.76 |
| k__Bacteria;p__Actinobacteria;c__Actinobacteria;o__Actinomycetales;f__ | 0 | 0 | 2 | 12 | 29 | 56.96 | 98.52 |
| k__Bacteria;p__Planctomycetes;c__Planctomycetia;o__Gemmatales;f__Gemmataceae | 0 | 0 | 2 | 12 | 24 | 33 | 42.88 |
| k__Bacteria;p__Acidobacteria;c__Solibacteres;o__Solibacterales;f__Solibacteraceae | 0 | 2 | 6 | 11 | 17 | 27 | 41.88 |
| k__Bacteria;p__Acidobacteria;c__TM1;o__;f__ | 0 | 0 | 0 | 11 | 49 | 97 | 149.76 |
| k__Bacteria;p__Acidobacteria;c__DA052;o__Ellin6513;f__ | 0 | 0 | 0 | 10 | 36 | 58 | 91.88 |
| k__Bacteria;p__Proteobacteria;c__Alphaproteobacteria;o__Rhizobiales;f__Bradyrhizobiaceae | 0 | 1 | 3 | 10 | 27 | 47 | 64 |
| k__Bacteria;p__Bacteroidetes;c__[Saprospirae];o__[Saprospirales];f__Chitinophagaceae | 0 | 0 | 2 | 9 | 22 | 60.96 | 113.88 |
| k__Bacteria;p__Chloroflexi;c__Ellin6529;o__;f__ | 0 | 0 | 3 | 9 | 17 | 31 | 45 |
| k__Bacteria;p__Actinobacteria;c__Actinobacteria;o__Actinomycetales;f__Streptomycetaceae | 0 | 3 | 5 | 9 | 14 | 20 | 41.88 |
| k__Bacteria;p__Actinobacteria;c__Thermoleophilia;o__Solirubrobacterales;f__ | 0 | 2 | 4 | 9 | 15 | 25 | 57.52 |
| k__Bacteria;p__Firmicutes;c__Bacilli;o__Bacillales;__ | 0 | 0 | 2 | 8 | 19 | 37 | 92.88 |
| k__Bacteria;p__Actinobacteria;c__Actinobacteria;o__Actinomycetales;f__Micrococcaceae | 0 | 0.04 | 2 | 7 | 16 | 39.96 | 105.88 |
| k__Bacteria;p__Acidobacteria;c__Solibacteres;o__Solibacterales;f__ | 0 | 1 | 3 | 7 | 12 | 18 | 26.88 |
| k__Bacteria;p__Planctomycetes;c__Planctomycetia;o__Gemmatales;f__Isosphaeraceae | 0 | 0 | 3 | 7 | 12 | 18.96 | 29.76 |
| k__Bacteria;p__Actinobacteria;c__MB-A2-108;o__0319-7L14;f__ | 0 | 0 | 2 | 7 | 16 | 28 | 37 |
| k__Bacteria;p__Nitrospirae;c__Nitrospira;o__Nitrospirales;f__0319-6A21 | 0 | 0 | 1 | 7 | 25 | 44 | 64.64 |
| k__Bacteria;p__Firmicutes;c__Bacilli;o__Bacillales;f__Bacillaceae | 0 | 0 | 2 | 7 | 17 | 26 | 41 |
| k__Bacteria;p__Nitrospirae;c__Nitrospira;o__Nitrospirales;f__Nitrospiraceae | 0 | 0 | 2 | 6 | 13 | 22 | 29 |
| k__Bacteria;p__Chloroflexi;c__TK10;o__B07_WMSP1;f__ | 0 | 1 | 2 | 6 | 14 | 33 | 70.88 |
| k__Bacteria;p__Proteobacteria;c__Gammaproteobacteria;o__Xanthomonadales;f__Sinobacteraceae | 0 | 0 | 2 | 6 | 14 | 24 | 43.88 |
| k__Bacteria;p__Planctomycetes;c__Phycisphaerae;o__WD2101;f__ | 0 | 0 | 2 | 6 | 21 | 58.96 | 99.64 |
| k__Bacteria;p__Armatimonadetes;c__0319-6E2;o__;f__ | 0 | 1 | 3 | 6 | 10 | 15 | 23 |
| k__Bacteria;p__Proteobacteria;c__Deltaproteobacteria;o__Myxococcales;f__ | 0 | 0 | 2 | 5 | 11 | 20.96 | 51.76 |
| k__Bacteria;p__Proteobacteria;c__Gammaproteobacteria;o__Xanthomonadales;f__Xanthomonadaceae | 0 | 0 | 2 | 5 | 11 | 22 | 41.88 |
| k__Bacteria;p__Proteobacteria;c__Betaproteobacteria;o__SC-I-84;f__ | 0 | 0 | 1 | 5 | 11 | 20 | 31.88 |
| k__Bacteria;p__Proteobacteria;c__Betaproteobacteria;o__Burkholderiales;f__Comamonadaceae | 0 | 0 | 1 | 5 | 10 | 16 | 31 |
| k__Bacteria;p__Planctomycetes;c__Planctomycetia;o__Pirellulales;f__Pirellulaceae | 0 | 0 | 1 | 4 | 9 | 16 | 22.88 |
| k__Bacteria;p__Proteobacteria;c__Alphaproteobacteria;o__Sphingomonadales;f__Sphingomonadaceae | 0 | 0 | 1 | 4 | 11 | 24.96 | 41.76 |
| k__Bacteria;__;__;__;__ | 0 | 1 | 2 | 4 | 7 | 12 | 26.76 |
| k__Bacteria;p__Firmicutes;c__Bacilli;o__Bacillales;f__Paenibacillaceae | 0 | 0 | 0 | 4 | 7 | 12 | 26.88 |
| k__Bacteria;p__Proteobacteria;c__Betaproteobacteria;o__MND1;f__ | 0 | 0 | 0 | 4 | 12 | 22 | 29.88 |
| k__Bacteria;p__Actinobacteria;c__Thermoleophilia;o__Solirubrobacterales;f__Conexibacteraceae | 0 | 0 | 1 | 4 | 7 | 9.96 | 12 |
| k__Bacteria;p__GAL15;c__;o__;f__ | 0 | 0 | 0 | 4 | 13 | 21 | 29 |
| k__Bacteria;p__Gemmatimonadetes;c__Gemmatimonadetes;o__;f__ | 0 | 0 | 1 | 4 | 12 | 19 | 31.76 |
| k__Bacteria;p__Actinobacteria;c__Acidimicrobiia;o__Acidimicrobiales;f__ | 0 | 0 | 2 | 4 | 7 | 9.96 | 13 |
| k__Bacteria;p__Actinobacteria;c__Actinobacteria;o__Actinomycetales;f__Micromonosporaceae | 0 | 0 | 1 | 3 | 10 | 31 | 52.88 |
| k__Bacteria;p__Actinobacteria;c__Actinobacteria;o__Actinomycetales;f__Nocardioidaceae | 0 | 0 | 0 | 3 | 8 | 17 | 35.88 |

| | | | | | | | | |
|---|---|---|---|---|---|---|---|---|
| k__Bacteria;p__Chloroflexi;c__Ktedonobacteria;o__Ktedonobacterales;f__Ktedonobacteraceae | | 0 | 0 | 0 | 3 | 7 | 13.96 | 21 |
| k__Bacteria;p__Proteobacteria;c__Betaproteobacteria;o__Burkholderiales;f__Oxalobacteraceae | | 0 | 0 | 1 | 3 | 8 | 17 | 33.88 |
| k__Bacteria;p__Verrucomicrobia;c__[Pedosphaerae];o__[Pedosphaerales];f__ | | 0 | 0 | 0 | 3 | 10 | 16 | 22 |
| k__Bacteria;p__Acidobacteria;c__iii1-8;o__DS-18;f__ | | 0 | 0 | 1 | 3 | 7 | 12.96 | 31 |
| k__Bacteria;p__Verrucomicrobia;c__[Pedosphaerae];o__[Pedosphaerales];f__Ellin515 | | 0 | 0 | 0 | 3 | 6 | 9 | 11.88 |
| k__Bacteria;p__Proteobacteria;c__Betaproteobacteria;o__;f__ | | 0 | 0 | 0 | 3 | 7 | 15 | 22 |
| k__Bacteria;p__Actinobacteria;c__Actinobacteria;o__Actinomycetales;f__Thermomonosporaceae | | 0 | 0 | 0 | 2 | 4 | 6 | 13.76 |
| k__Bacteria;p__Acidobacteria;c__[Chloracidobacteria];o__11-24;f__ | | 0 | 0 | 0 | 2 | 5 | 10 | 17.64 |
| k__Bacteria;p__Actinobacteria;c__Actinobacteria;o__Actinomycetales;f__Pseudonocardiaceae | | 0 | 0 | 0 | 2 | 5 | 9 | 15 |
| k__Bacteria;p__Proteobacteria;c__Alphaproteobacteria;o__Rhodospirillales;f__Acetobacteraceae | | 0 | 0 | 1 | 2 | 5 | 8 | 16 |
| k__Bacteria;p__Gemmatimonadetes;c__Gemmatimonadetes;o__N1423WL;f__ | | 0 | 0 | 0 | 2 | 6 | 11 | 21.76 |
| k__Bacteria;p__Proteobacteria;c__Betaproteobacteria;o__Burkholderiales;f__Burkholderiaceae | | 0 | 0 | 0 | 2 | 5 | 11.96 | 56.76 |
| k__Bacteria;p__Proteobacteria;c__Deltaproteobacteria;o__Myxococcales;f__Haliangiaceae | | 0 | 0 | 0 | 2 | 5 | 9 | 15.88 |
| k__Bacteria;p__Planctomycetes;c__Planctomycetia;o__Planctomycetales;f__Planctomycetaceae | | 0 | 0 | 0 | 2 | 3 | 6 | 9.88 |
| k__Bacteria;p__Actinobacteria;c__Acidimicrobiia;o__Acidimicrobiales;f__EB1017 | | 0 | 0 | 0 | 2 | 4 | 7 | 11 |
| k__Bacteria;p__Proteobacteria;__;__;__ | | 0 | 0 | 0 | 2 | 5 | 9 | 13.88 |
| k__Bacteria;p__Chloroflexi;c__S085;o__;f__ | | 0 | 0 | 0 | 2 | 5 | 9 | 13 |
| k__Bacteria;p__Proteobacteria;c__Alphaproteobacteria;o__Ellin329;f__ | | 0 | 0 | 0 | 2 | 4 | 10 | 18.88 |
| k__Bacteria;p__Proteobacteria;c__Alphaproteobacteria;o__Rhizobiales;__ | | 0 | 0 | 0 | 2 | 4 | 10 | 16 |
| k__Bacteria;p__Proteobacteria;c__Betaproteobacteria;o__Ellin6067;f__ | | 0 | 0 | 0 | 2 | 5 | 9 | 15.88 |
| k__Bacteria;p__Proteobacteria;c__Gammaproteobacteria;o__Legionellales;f__Coxiellaceae | | 0 | 0 | 0 | 1 | 2 | 3.96 | 6 |
| k__Bacteria;p__Chloroflexi;c__Ktedonobacteria;o__TK10;f__ | | 0 | 0 | 0 | 1 | 5 | 11 | 18 |
| k__Bacteria;p__Acidobacteria;c__Sva0725;o__Sva0725;f__ | | 0 | 0 | 0 | 1 | 4 | 9 | 21 |
| k__Bacteria;p__Proteobacteria;c__Deltaproteobacteria;o__Myxococcales;__ | | 0 | 0 | 0 | 1 | 4 | 13 | 30.88 |
| k__Bacteria;p__WPS-2;c__;o__;f__ | | 0 | 0 | 0 | 1 | 7 | 14 | 23.76 |
| k__Bacteria;p__Chloroflexi;c__Ktedonobacteria;o__JG30-KF-AS9;f__ | | 0 | 0 | 0 | 1 | 4 | 8 | 16 |
| k__Bacteria;p__Proteobacteria;c__Alphaproteobacteria;o__Caulobacterales;f__Caulobacteraceae | | 0 | 0 | 0 | 1 | 2 | 4 | 7 |
| k__Bacteria;p__Chloroflexi;__;__;__ | | 0 | 0 | 0 | 1 | 4 | 10.96 | 16 |
| k__Bacteria;p__Chloroflexi;c__TK10;o__B07_WMSP1;f__FFCH4570 | | 0 | 0 | 0 | 1 | 2 | 4 | 7.88 |
| k__Bacteria;p__Actinobacteria;c__Thermoleophilia;o__Gaiellales;f__ | | 0 | 0 | 0 | 1 | 2 | 5 | 8 |
| k__Bacteria;p__Acidobacteria;c__Acidobacteriia;o__Acidobacteriales;f__Acidobacteriaceae | | 0 | 0 | 0 | 1 | 4 | 10.96 | 21 |
| k__Bacteria;p__Firmicutes;c__Bacilli;o__Bacillales;f__Planococcaceae | | 0 | 0 | 0 | 1 | 3 | 9 | 24.88 |
| k__Archaea;p__Crenarchaeota;c__Thaumarchaeota;o__Cenarchaeales;f__SAGMA-X | | 0 | 0 | 0 | 1 | 4 | 10 | 24.88 |
| k__Bacteria;p__Proteobacteria;c__Alphaproteobacteria;o__Rhizobiales;f__ | | 0 | 0 | 0 | 1 | 4 | 9 | 14 |
| k__Bacteria;p__Actinobacteria;c__MB-A2-108;o__;f__ | | 0 | 0 | 0 | 1 | 2 | 5 | 9 |
| k__Bacteria;p__Chloroflexi;c__Ktedonobacteria;o__B12-WMSP1;f__ | | 0 | 0 | 0 | 1 | 8 | 21.96 | 40 |
| k__Bacteria;p__Armatimonadetes;c__Chthonomonadetes;o__SJA-22;f__ | | 0 | 0 | 0 | 1 | 2 | 3 | 5 |
| k__Bacteria;p__Proteobacteria;c__Deltaproteobacteria;o__Myxococcales;f__Myxococcaceae | | 0 | 0 | 0 | 1 | 3 | 5 | 9 |
| k__Bacteria;p__Actinobacteria;c__Actinobacteria;o__Actinomycetales;f__Intrasporangiaceae | | 0 | 0 | 0 | 1 | 3 | 7.96 | 14 |
| k__Bacteria;p__Acidobacteria;c__Acidobacteria-6;o__iii1-15;f__mb2424 | | 0 | 0 | 0 | 1 | 5 | 10 | 22.88 |
| k__Bacteria;p__Acidobacteria;c__Acidobacteria-5;o__;f__ | | 0 | 0 | 0 | 1 | 3 | 7 | 10 |
| k__Bacteria;p__Actinobacteria;c__Actinobacteria;o__Actinomycetales;f__Geodermatophilaceae | | 0 | 0 | 0 | 1 | 4 | 11 | 20 |
| k__Bacteria;p__Chloroflexi;c__C0119;o__;f__ | | 0 | 0 | 0 | 1 | 3 | 5 | 9 |
| k__Bacteria;p__Acidobacteria;c__Acidobacteria-6;o__iii1-15;f__RB40 | | 0 | 0 | 0 | 1 | 4 | 10 | 15.88 |
| k__Bacteria;p__WS3;c__PRR-12;o__Sediment-1;f__ | | 0 | 0 | 0 | 1 | 5 | 15 | 25.76 |
| k__Bacteria;p__Proteobacteria;c__Zetaproteobacteria;o__Mariprofundales;f__Mariprofundaceae | | 0 | 0 | 0 | 1 | 3 | 8 | 11 |
| k__Bacteria;p__Acidobacteria;c__Acidobacteria-6;o__CCU21;f__ | | 0 | 0 | 0 | 1 | 5 | 10.96 | 16.88 |
| k__Bacteria;p__Proteobacteria;c__Alphaproteobacteria;o__Rhizobiales;f__Phyllobacteriaceae | | 0 | 0 | 0 | 1 | 3 | 6 | 8 |
| k__Bacteria;p__Bacteroidetes;c__Cytophagia;o__Cytophagales;f__Cytophagaceae | | 0 | 0 | 0 | 1 | 3 | 9.96 | 23 |
| k__Bacteria;p__Actinobacteria;c__Actinobacteria;o__Actinomycetales;__ | | 0 | 0 | 0 | 1 | 3 | 8 | 20 |

| | | | | | | | |
|---|---|---|---|---|---|---|---|
| k__Bacteria;p__Actinobacteria;c__Actinobacteria;o__Actinomycetales;f__Mycobacteriaceae | 0 | 0 | 0 | 1 | 3 | 6 | 9.88 |
| k__Bacteria;p__Actinobacteria;c__Thermoleophilia;o__Solirubrobacterales;f__Solirubrobacteraceae | 0 | 0 | 0 | 1 | 5 | 12 | 20.88 |
| k__Bacteria;p__Acidobacteria;c__iii1-8;o__32-20;f__ | 0 | 0 | 0 | 1 | 4 | 7 | 10 |
| k__Bacteria;p__AD3;c__JG37-AG-4;o__;f__ | 0 | 0 | 0 | 1 | 7 | 18.96 | 35.88 |
| k__Bacteria;p__Gemmatimonadetes;c__Gemmatimonadetes;o__Ellin5290;f__ | 0 | 0 | 0 | 1 | 3 | 8 | 14.88 |
| k__Bacteria;p__Actinobacteria;c__Actinobacteria;o__Actinomycetales;f__Streptosporangiaceae | 0 | 0 | 0 | 1 | 2 | 4 | 9.88 |
| k__Bacteria;p__Acidobacteria;c__[Chloracidobacteria];o__RB41;f__Ellin6075 | 0 | 0 | 0 | 1 | 5 | 12 | 24.88 |
| k__Archaea;p__Crenarchaeota;c__MBGA;o__NRP-J;f__ | 0 | 0 | 0 | 1 | 7 | 13 | 25.76 |

| Feature ID | 2% | 9% | 25% | 50% | 75% | 91% | 98% |
|---|---|---|---|---|---|---|---|
| k__Bacteria;p__Acidobacteria;c__Acidobacteria-6;o__iii1-15;f__ | 13.5 | 30 | 47 | 68 | 90 | 109.25 | 126 |
| k__Archaea;p__Crenarchaeota;c__Thaumarchaeota;o__Nitrososphaerales;f__Nitrososphaeraceae | 7 | 19.75 | 34.75 | 48 | 67 | 94.25 | 145 |
| k__Bacteria;p__Actinobacteria;c__Thermoleophilia;o__Gaiellales;f__Gaiellaceae | 3 | 16.75 | 29.75 | 42 | 58.25 | 81 | 111.5 |
| k__Bacteria;p__Acidobacteria;c__[Chloracidobacteria];o__RB41;f__ | 2 | 13 | 24 | 38 | 54 | 73.5 | 103.5 |
| k__Bacteria;p__Proteobacteria;c__Alphaproteobacteria;o__Rhizobiales;f__Hyphomicrobiaceae | 0 | 2 | 16 | 30 | 43 | 53.25 | 64.5 |
| k__Bacteria;p__Proteobacteria;c__Deltaproteobacteria;o__Syntrophobacterales;f__Syntrophobacteraceae | 0 | 2 | 10.75 | 30 | 45 | 53 | 62 |
| k__Bacteria;p__Verrucomicrobia;c__[Spartobacteria];o__[Chthoniobacterales];f__[Chthoniobacteraceae] | 0 | 0 | 12 | 29.5 | 49.25 | 84 | 143.5 |
| k__Bacteria;p__Proteobacteria;c__Alphaproteobacteria;o__Rhodospirillales;f__Rhodospirillaceae | 6 | 11 | 17 | 27 | 39 | 49 | 54 |
| k__Bacteria;p__Proteobacteria;c__Alphaproteobacteria;o__Rhizobiales;f__Bradyrhizobiaceae | 0.5 | 3 | 9 | 26 | 41 | 58.25 | 76 |
| k__Bacteria;p__Planctomycetes;c__Phycisphaerae;o__WD2101;f__ | 0 | 3 | 7 | 21 | 49.25 | 76.25 | 115 |
| k__Bacteria;p__Bacteroidetes;c__[Saprospirae];o__[Saprospirales];f__Chitinophagaceae | 2 | 6 | 11.75 | 20 | 49.25 | 92.25 | 123.5 |
| k__Bacteria;p__Acidobacteria;c__Acidobacteriia;o__Acidobacteriales;f__Koribacteraceae | 0 | 3 | 9 | 19 | 45 | 85 | 118.5 |
| k__Bacteria;p__Firmicutes;c__Bacilli;o__Bacillales;__ | 2 | 4 | 11 | 18 | 33 | 59.5 | 119 |
| k__Bacteria;p__Planctomycetes;c__Planctomycetia;o__Gemmatales;f__Gemmataceae | 0 | 0 | 0 | 18 | 28 | 35.25 | 43.5 |
| k__Bacteria;p__Gemmatimonadetes;c__Gemm-1;o__;f__ | 1 | 4 | 8 | 17 | 39 | 53.5 | 65.5 |
| k__Bacteria;p__Actinobacteria;c__Thermoleophilia;o__Solirubrobacterales;f__ | 1 | 4 | 9 | 15 | 21 | 40.25 | 69 |
| k__Bacteria;p__Chloroflexi;c__Ellin6529;o__;f__ | 1 | 4 | 8 | 14 | 24 | 34 | 45.5 |
| k__Bacteria;p__Proteobacteria;c__Gammaproteobacteria;o__Xanthomonadales;f__Sinobacteraceae | 0 | 3 | 8 | 14 | 21 | 31 | 51 |
| k__Bacteria;p__Firmicutes;c__Bacilli;o__Bacillales;f__Bacillaceae | 2 | 4.75 | 8 | 14 | 23 | 31.25 | 47 |
| k__Bacteria;p__Proteobacteria;c__Deltaproteobacteria;o__Myxococcales;f__ | 1 | 4 | 7 | 11 | 17 | 34 | 59 |
| k__Bacteria;p__Gemmatimonadetes;c__Gemmatimonadetes;o__;f__ | 0.5 | 2.75 | 6.75 | 11 | 16 | 25 | 38.5 |
| k__Bacteria;p__Nitrospirae;c__Nitrospira;o__Nitrospirales;f__0319-6A21 | 0 | 0 | 1 | 11 | 34 | 47.25 | 56.5 |
| k__Bacteria;p__Acidobacteria;c__Solibacteres;o__Solibacterales;f__Solibacteraceae | 0 | 1 | 4 | 11 | 19 | 30 | 47 |
| k__Bacteria;p__Proteobacteria;c__Betaproteobacteria;o__MND1;f__ | 0 | 0.75 | 5 | 10 | 19 | 24.25 | 31 |
| k__Bacteria;p__Actinobacteria;c__Actinobacteria;o__Actinomycetales;f__Streptomycetaceae | 0.5 | 3 | 5 | 9 | 13 | 17 | 29.5 |
| k__Bacteria;p__Acidobacteria;c__Solibacteres;o__Solibacterales;f__ | 0 | 2 | 5 | 9 | 16 | 22 | 30.5 |
| k__Bacteria;p__Proteobacteria;c__Betaproteobacteria;o__Burkholderiales;f__Comamonadaceae | 1 | 2 | 4 | 8.5 | 14 | 19.25 | 34 |
| k__Bacteria;p__Proteobacteria;c__Betaproteobacteria;o__SC-I-84;f__ | 0 | 2 | 4 | 8 | 17 | 24.25 | 34.5 |
| k__Bacteria;p__Nitrospirae;c__Nitrospira;o__Nitrospirales;f__Nitrospiraceae | 0 | 0 | 2 | 8 | 17.25 | 25 | 31 |
| k__Bacteria;p__Verrucomicrobia;c__[Pedosphaerae];o__[Pedosphaerales];f__ | 0 | 0 | 0 | 8 | 14 | 18 | 23 |
| k__Bacteria;p__Actinobacteria;c__Actinobacteria;o__Actinomycetales;f__Micromonosporaceae | 0 | 1 | 3.75 | 8 | 22.25 | 44.25 | 62.5 |
| k__Bacteria;p__Planctomycetes;c__Planctomycetia;o__Pirellulales;f__Pirellulaceae | 0 | 0 | 4 | 8 | 13 | 19.25 | 26 |
| k__Bacteria;p__Actinobacteria;c__MB-A2-108;o__0319-7L14;f__ | 0 | 0 | 1 | 8 | 21 | 33 | 39 |
| k__Bacteria;p__Proteobacteria;c__Gammaproteobacteria;o__Xanthomonadales;f__Xanthomonadaceae | 0 | 0.75 | 2 | 7 | 16 | 31 | 54 |
| k__Bacteria;p__Actinobacteria;c__Actinobacteria;o__Actinomycetales;f__Micrococcaceae | 0 | 1 | 2 | 7 | 13.25 | 32.25 | 57.5 |
| k__Bacteria;p__Planctomycetes;c__Planctomycetia;o__Gemmatales;f__Isosphaeraceae | 0 | 0 | 0 | 6 | 10 | 17 | 23.5 |
| k__Bacteria;p__Proteobacteria;c__Betaproteobacteria;o__;f__ | 0 | 1 | 2 | 6 | 13 | 19 | 24.5 |
| k__Bacteria;p__Actinobacteria;c__Actinobacteria;o__Actinomycetales;f__Nocardioidaceae | 0 | 1 | 2 | 6 | 12 | 25 | 43.5 |
| k__Bacteria;p__Acidobacteria;c__iii1-8;o__DS-18;f__ | 1 | 2 | 3 | 6 | 10 | 18.25 | 38 |
| k__Bacteria;p__Proteobacteria;c__Betaproteobacteria;o__Burkholderiales;f__Oxalobacteraceae | 0 | 1 | 2 | 5 | 13.25 | 23 | 35 |
| k__Bacteria;p__Proteobacteria;c__Betaproteobacteria;o__Ellin6067;f__ | 0 | 1 | 2 | 5 | 8 | 13 | 19 |
| k__Bacteria;p__Gemmatimonadetes;c__Gemmatimonadetes;o__N1423WL;f__ | 0 | 0 | 2 | 5 | 9 | 15.25 | 26 |
| k__Bacteria;p__Proteobacteria;c__Alphaproteobacteria;o__Sphingomonadales;f__Sphingomonadaceae | 0 | 1 | 2.75 | 5 | 15 | 28 | 42.5 |
| k__Bacteria;p__Chloroflexi;c__TK10;o__B07_WMSP1;f__ | 0 | 0 | 2 | 5 | 10.25 | 19.25 | 31.5 |
| k__Bacteria;p__Acidobacteria;c__Acidobacteria-6;o__iii1-15;f__mb2424 | 0 | 0 | 2 | 4.5 | 8 | 15 | 26 |
| k__Bacteria;p__Actinobacteria;c__Acidimicrobiia;o__Acidimicrobiales;f__ | 0 | 0 | 0 | 4 | 7 | 10 | 13.5 |

| | | | | | | | |
|---|---|---|---|---|---|---|---|
| k__Bacteria;p__Proteobacteria;c__Deltaproteobacteria;o__Myxococcales;__ | 0 | 0.75 | 2 | 4 | 10 | 22.25 | 35.5 |
| k__Bacteria;p__Armatimonadetes;c__0319-6E2;o__;f__ | 0 | 0 | 2 | 4 | 7 | 10 | 13.5 |
| k__Bacteria;p__Actinobacteria;c__Actinobacteria;o__Actinomycetales;f__Pseudonocardiaceae | 0 | 1 | 2 | 4 | 8 | 11.25 | 16.5 |
| k__Bacteria;p__Acidobacteria;c__[Chloracidobacteria];o__11-24;f__ | 0 | 0 | 1 | 4 | 7 | 10 | 14.5 |
| k__Bacteria;p__Acidobacteria;c__[Chloracidobacteria];o__RB41;f__Ellin6075 | 0 | 0 | 1 | 4 | 9 | 18 | 35.5 |
| k__Bacteria;p__Acidobacteria;c__Sva0725;o__Sva0725;f__ | 0 | 0 | 2 | 4 | 7 | 13 | 27 |
| k__Bacteria;p__Proteobacteria;c__Deltaproteobacteria;o__Myxococcales;f__Haliangiaceae | 0 | 1 | 3 | 4 | 8 | 12 | 18.5 |
| k__Bacteria;p__Proteobacteria;c__Alphaproteobacteria;o__Rhizobiales;__ | 0 | 0 | 2 | 3.5 | 8 | 13 | 18 |
| k__Bacteria;p__Proteobacteria;c__Alphaproteobacteria;o__Rhodospirillales;f__Acetobacteraceae | 0 | 0 | 0.75 | 3 | 6 | 12 | 19 |
| k__Bacteria;p__Actinobacteria;c__Actinobacteria;o__Actinomycetales;f__Geodermatophilaceae | 0 | 0 | 0 | 3 | 7 | 13.25 | 24 |
| k__Bacteria;p__Verrucomicrobia;c__[Pedosphaerae];o__[Pedosphaerales];f__Ellin515 | 0 | 0 | 0 | 3 | 7 | 10 | 13 |
| k__Bacteria;p__Acidobacteria;c__iii1-8;o__32-20;f__ | 0 | 0 | 0 | 3 | 5 | 8 | 11 |
| k__Bacteria;__;__;__;__ | 0 | 1 | 2 | 3 | 5 | 10 | 356.5 |
| k__Bacteria;p__Actinobacteria;c__Thermoleophilia;o__Solirubrobacterales;f__Solirubrobacteraceae | 0 | 0 | 1 | 3 | 9 | 15 | 25 |
| k__Bacteria;p__Firmicutes;c__Bacilli;o__Bacillales;f__Paenibacillaceae | 0 | 0 | 0 | 3 | 7 | 12 | 15.5 |
| k__Bacteria;p__Proteobacteria;c__Zetaproteobacteria;o__Mariprofundales;f__Mariprofundaceae | 0 | 0 | 0 | 2.5 | 6 | 9 | 13 |
| k__Bacteria;p__Acidobacteria;c__Acidobacteria-6;o__CCU21;f__ | 0 | 0 | 0 | 2.5 | 7 | 13 | 18 |
| k__Bacteria;p__Acidobacteria;c__Acidobacteria-5;o__;f__ | 0 | 0 | 0 | 2 | 5 | 8 | 11.5 |
| k__Bacteria;p__Actinobacteria;c__Thermoleophilia;o__Solirubrobacterales;f__Conexibacteraceae | 0 | 0 | 0 | 2 | 5 | 9 | 12 |
| k__Bacteria;p__Gemmatimonadetes;c__Gemmatimonadetes;o__Ellin5290;f__ | 0 | 0 | 0 | 2 | 5 | 10.25 | 17.5 |
| k__Bacteria;p__Chloroflexi;c__Thermomicrobia;o__JG30-KF-CM45;f__ | 0 | 0 | 0 | 2 | 7 | 14.25 | 26.5 |
| k__Bacteria;p__WS3;c__PRR-12;o__Sediment-1;f__PRR-10 | 0 | 0 | 0 | 2 | 6 | 11 | 17 |
| k__Bacteria;p__Actinobacteria;c__Actinobacteria;o__Actinomycetales;f__ | 0 | 0 | 0 | 2 | 10.25 | 17 | 28 |
| k__Bacteria;p__Actinobacteria;c__Actinobacteria;o__Actinomycetales;f__Thermomonosporaceae | 0 | 0 | 0 | 2 | 4 | 6 | 9 |
| k__Bacteria;p__Gemmatimonadetes;c__Gemmatimonadetes;o__Gemmatimonadales;f__Ellin5301 | 0 | 0 | 0 | 2 | 5 | 8 | 11 |
| k__Bacteria;p__Actinobacteria;c__Actinobacteria;o__Actinomycetales;f__Mycobacteriaceae | 0 | 0 | 0 | 2 | 4 | 6.25 | 12 |
| k__Bacteria;p__Proteobacteria;c__Alphaproteobacteria;o__Rhizobiales;f__ | 0 | 0 | 0 | 2 | 6.25 | 11 | 18 |
| k__Bacteria;p__Chloroflexi;c__S085;o__;f__ | 0 | 0 | 0 | 2 | 6 | 10 | 14 |
| k__Bacteria;p__Proteobacteria;c__Deltaproteobacteria;o__Myxococcales;f__Myxococcaceae | 0 | 0 | 0 | 2 | 4 | 6 | 11 |
| k__Bacteria;p__Planctomycetes;c__Planctomycetia;o__Planctomycetales;f__Planctomycetaceae | 0 | 0 | 0 | 2 | 4 | 8 | 10.5 |
| k__Bacteria;p__Acidobacteria;c__Acidobacteria-6;o__iii1-15;f__RB40 | 0 | 0 | 0 | 2 | 8 | 13.25 | 18 |
| k__Bacteria;p__WS3;c__PRR-12;o__Sediment-1;f__ | 0 | 0 | 0 | 2 | 11 | 21 | 31.5 |
| k__Bacteria;p__Proteobacteria;c__Alphaproteobacteria;o__Ellin329;f__ | 0 | 0 | 0 | 2 | 7 | 13 | 26.5 |
| k__Bacteria;p__Actinobacteria;c__Actinobacteria;o__Actinomycetales;__ | 0 | 0 | 1 | 2 | 6 | 14.25 | 26.5 |
| k__Bacteria;p__Proteobacteria;c__Alphaproteobacteria;o__Rhizobiales;f__Phyllobacteriaceae | 0 | 0 | 0 | 2 | 4 | 6.25 | 9 |
| k__Bacteria;p__Proteobacteria;c__Deltaproteobacteria;o__Desulfuromonadales;f__Geobacteraceae | 0 | 0 | 0 | 2 | 9 | 15 | 24.5 |
| k__Bacteria;p__Actinobacteria;c__Actinobacteria;o__Actinomycetales;f__Streptosporangiaceae | 0 | 0 | 0 | 2 | 3 | 5.25 | 15.5 |
| k__Bacteria;p__Verrucomicrobia;c__[Pedosphaerae];o__[Pedosphaerales];f__Ellin517 | 0 | 0 | 0 | 2 | 4 | 8 | 13 |
| k__Bacteria;p__Bacteroidetes;c__Cytophagia;o__Cytophagales;f__Cytophagaceae | 0 | 0 | 0 | 2 | 5 | 15 | 38 |
| k__Bacteria;p__Actinobacteria;c__Acidimicrobiia;o__Acidimicrobiales;f__EB1017 | 0 | 0 | 0 | 1.5 | 4 | 6 | 8 |
| k__Bacteria;p__Actinobacteria;c__Actinobacteria;o__Actinomycetales;f__Frankiaceae | 0 | 0 | 0 | 1 | 3 | 5.25 | 9.5 |
| k__Bacteria;p__Proteobacteria;c__Betaproteobacteria;o__Burkholderiales;f__Burkholderiaceae | 0 | 0 | 0 | 1 | 3 | 8 | 16 |
| k__Bacteria;p__Actinobacteria;c__Thermoleophilia;o__Gaiellales;f__ | 0 | 0 | 0 | 1 | 4 | 6 | 9 |
| k__Bacteria;p__Chloroflexi;c__Ktedonobacteria;o__Thermogemmatisporales;f__Thermogemmatisporaceae | 0 | 0 | 0 | 1 | 7 | 40 | 133 |
| k__Bacteria;p__Acidobacteria;c__[Chloracidobacteria];o__PK29;f__ | 0 | 0 | 0 | 1 | 2 | 4 | 6 |
| k__Bacteria;p__Actinobacteria;c__Thermoleophilia;o__Solirubrobacterales;f__Patulibacteraceae | 0 | 0 | 0 | 1 | 3 | 6 | 12 |
| k__Bacteria;p__Actinobacteria;c__Actinobacteria;o__Actinomycetales;f__Intrasporangiaceae | 0 | 0 | 0 | 1 | 3 | 5 | 11 |

| | | | | | | | |
|---|---|---|---|---|---|---|---|
| k__Bacteria;p__Chloroflexi;c__Ktedonobacteria;o__Ktedonobacterales;f__Ktedonobacteraceae | 0 | 0 | 0 | 1 | 3 | 6.25 | 12.5 |
| k__Bacteria;p__Chloroflexi;c__Gitt-GS-136;o__;f__ | 0 | 0 | 0 | 1 | 4 | 7.25 | 10 |
| k__Bacteria;p__Armatimonadetes;c__Chthonomonadetes;o__SJA-22;f__ | 0 | 0 | 0 | 1 | 2 | 4 | 6 |
| k__Bacteria;p__Firmicutes;c__Bacilli;o__Bacillales;f__Planococcaceae | 0 | 0 | 0 | 1 | 3 | 14.25 | 38.5 |
| k__Bacteria;p__Chloroflexi;c__Anaerolineae;o__SBR1031;f__A4b | 0 | 0 | 0 | 1 | 3 | 6 | 9 |
| k__Bacteria;p__Actinobacteria;c__Rubrobacteria;o__Rubrobacterales;f__Rubrobacteraceae | 0 | 0 | 0 | 1 | 8 | 81.25 | 230.5 |
| k__Bacteria;p__Acidobacteria;c__S035;o__;f__ | 0 | 0 | 0 | 1 | 7 | 10 | 13 |
| k__Bacteria;p__Chloroflexi;c__Anaerolineae;o__SBR1031;f__oc28 | 0 | 0 | 0 | 1 | 2 | 4 | 6.5 |
| k__Bacteria;p__Actinobacteria;c__MB-A2-108;o__;f__ | 0 | 0 | 0 | 1 | 3 | 6 | 9 |
| k__Bacteria;p__;c__;o__;f__ | 0 | 0 | 0 | 1 | 2 | 4 | 6 |
| k__Bacteria;p__Verrucomicrobia;c__Opitutae;o__Opitutales;f__Opitutaceae | 0 | 0 | 0 | 1 | 3 | 9 | 20 |
| k__Bacteria;p__Proteobacteria;c__Deltaproteobacteria;o__Myxococcales;f__Cystobacteraceae | 0 | 0 | 0 | 1 | 3 | 7.25 | 18.5 |
| k__Bacteria;p__Proteobacteria;c__Alphaproteobacteria;o__Rhizobiales;f__Beijerinckiaceae | 0 | 0 | 0 | 1 | 6 | 21 | 28.5 |
| k__Bacteria;p__Proteobacteria;c__Alphaproteobacteria;o__Caulobacterales;f__Caulobacteraceae | 0 | 0 | 0 | 1 | 3 | 5 | 8.5 |
| k__Bacteria;p__Proteobacteria;c__Alphaproteobacteria;o__Rhizobiales;f__Rhizobiaceae | 0 | 0 | 0 | 1 | 3 | 7 | 13 |
| k__Bacteria;p__Proteobacteria;c__Betaproteobacteria;o__Rhodocyclales;f__Rhodocyclaceae | 0 | 0 | 0 | 1 | 4 | 8.25 | 17.5 |
| k__Bacteria;p__Actinobacteria;c__Actinobacteria;o__Actinomycetales;f__Actinosynnemataceae | 0 | 0 | 0 | 1 | 2.25 | 6 | 18 |
| k__Bacteria;p__Proteobacteria;c__Deltaproteobacteria;o__Myxococcales;f__Polyangiaceae | 0 | 0 | 0 | 1 | 2 | 4.25 | 11 |
| k__Bacteria;p__Acidobacteria;c__Acidobacteria-6;o__;f__ | 0 | 0 | 0 | 1 | 6.25 | 14 | 20 |
| k__Bacteria;p__Proteobacteria;c__Alphaproteobacteria;o__Rhizobiales;f__Xanthobacteraceae | 0 | 0 | 0 | 1 | 5 | 12 | 17.5 |
| k__Bacteria;p__Chloroflexi;c__Chloroflexi;o__[Roseiflexales];f__[Kouleothrixaceae] | 0 | 0 | 0 | 1 | 2 | 5 | 7 |
| k__Bacteria;p__Acidobacteria;c__Acidobacteria-6;__;__ | 0 | 0 | 0 | 1 | 7 | 11 | 16 |
| k__Bacteria;p__Proteobacteria;c__Deltaproteobacteria;o__[Entotheonellales];f__[Entotheonellaceae] | 0 | 0 | 0 | 1 | 4 | 22 | 34.5 |
| k__Bacteria;p__Chloroflexi;c__TK10;o__;f__ | 0 | 0 | 0 | 1 | 2 | 9 | 17.5 |
| k__Bacteria;p__Actinobacteria;c__Thermoleophilia;o__Solirubrobacterales;__ | 0 | 0 | 0 | 1 | 2 | 6.25 | 10 |
| k__Bacteria;p__Chloroflexi;c__C0119;o__;f__ | 0 | 0 | 0 | 1 | 3 | 6 | 10.5 |
| k__Bacteria;p__Actinobacteria;c__Actinobacteria;o__Actinomycetales;f__Kineosporiaceae | 0 | 0 | 0 | 1 | 3 | 8 | 12 |
| k__Bacteria;p__Chloroflexi;c__Anaerolineae;o__envOPS12;f__ | 0 | 0 | 0 | 0.5 | 4 | 27 | 76 |
| k__Bacteria;p__Proteobacteria;c__Betaproteobacteria;o__A21b;f__EB1003 | 0 | 0 | 0 | 0.5 | 2 | 5 | 7 |

| Feature ID | 2% | 9% | 25% | 50% | 75% | 91% | 98% |
|---|---|---|---|---|---|---|---|
| k__Bacteria;p__Chloroflexi;c__Ktedonobacteria;o__Thermogemmatisporales;f__Thermogemmatisporaceae | 54.6 | 94.4 | 148 | 223 | 309 | 414.4 | 592.6 |
| k__Bacteria;p__AD3;c__ABS-6;o__;f__ | 17.6 | 29.2 | 70 | 117 | 203 | 287 | 400 |
| k__Bacteria;p__Acidobacteria;c__Acidobacteriia;o__Acidobacteriales;f__Koribacteraceae | 27.6 | 40 | 55 | 77 | 113 | 145 | 174.4 |
| k__Bacteria;p__Acidobacteria;c__TM1;o__;f__ | 7 | 13 | 24 | 48 | 82 | 122.8 | 162.8 |
| k__Bacteria;p__Verrucomicrobia;c__[Spartobacteria];o__[Chthoniobacterales];f__[Chthoniobacteraceae] | 4.6 | 12 | 24 | 36 | 56 | 88 | 125 |
| k__Bacteria;p__Acidobacteria;c__DA052;o__Ellin6513;f__ | 5.6 | 13 | 23 | 36 | 53 | 71.8 | 108 |
| k__Bacteria;p__Actinobacteria;c__Thermoleophilia;o__Gaiellales;f__Gaiellaceae | 11 | 17.2 | 24 | 33 | 46 | 63 | 78.8 |
| k__Bacteria;p__Proteobacteria;c__Alphaproteobacteria;o__Rhodospirillales;f__Rhodospirillaceae | 8.6 | 14 | 20 | 28 | 37 | 53.8 | 75.4 |
| k__Bacteria;p__Actinobacteria;c__Actinobacteria;o__Actinomycetales;f__ | 4 | 7 | 13 | 27 | 47 | 72.8 | 106 |
| k__Bacteria;p__Proteobacteria;c__Deltaproteobacteria;o__Syntrophobacterales;f__Syntrophobacteraceae | 6 | 10.2 | 16 | 22 | 30 | 39 | 54.4 |
| k__Bacteria;p__Proteobacteria;c__Alphaproteobacteria;o__Rhizobiales;f__Hyphomicrobiaceae | 6.6 | 9 | 13 | 20 | 29 | 41 | 57 |
| k__Bacteria;p__Acidobacteria;c__Acidobacteria-6;o__iii1-15;f__ | 0 | 2 | 5 | 16 | 32 | 54 | 70 |
| k__Bacteria;p__Gemmatimonadetes;c__Gemm-1;o__;f__ | 0.6 | 3 | 7 | 14 | 21 | 37.8 | 62.6 |
| k__Archaea;p__Crenarchaeota;c__Thaumarchaeota;o__Nitrososphaerales;f__Nitrososphaeraceae | 1 | 3 | 7 | 13 | 22 | 31.8 | 42 |
| k__Bacteria;p__GAL15;c__;o__;f__ | 3 | 5 | 8 | 12 | 18 | 25 | 38.4 |
| k__Bacteria;p__Acidobacteria;c__Solibacteres;o__Solibacterales;f__Solibacteraceae | 1.6 | 4.2 | 8 | 12 | 16 | 22 | 32 |
| k__Bacteria;p__Actinobacteria;c__Actinobacteria;o__Actinomycetales;f__Streptomycetaceae | 0 | 2 | 6 | 10 | 15 | 23 | 57.4 |
| k__Bacteria;p__Planctomycetes;c__Planctomycetia;o__Gemmatales;f__Gemmataceae | 1 | 2 | 5 | 10 | 18 | 28.8 | 42.4 |
| k__Bacteria;p__Planctomycetes;c__Planctomycetia;o__Gemmatales;f__Isosphaeraceae | 1 | 2 | 5 | 9 | 14 | 19 | 40 |
| k__Bacteria;p__Armatimonadetes;c__0319-6E2;o__;f__ | 1 | 3 | 5 | 8 | 12 | 18 | 29 |
| k__Bacteria;p__Chloroflexi;c__TK10;o__B07_WMSP1;f__ | 0 | 1 | 4 | 8 | 23 | 45 | 88.6 |
| k__Bacteria;p__Chloroflexi;c__Ktedonobacteria;o__B12-WMSP1;f__ | 0 | 0 | 3 | 8 | 17 | 29.8 | 51.4 |
| k__Bacteria;p__Acidobacteria;c__[Chloracidobacteria];o__RB41;f__ | 0 | 0 | 2 | 7 | 16 | 28 | 40.4 |
| k__Bacteria;p__Actinobacteria;c__MB-A2-108;o__0319-7L14;f__ | 0 | 1 | 3 | 7 | 13 | 22 | 32 |
| k__Bacteria;p__Actinobacteria;c__Actinobacteria;o__Actinomycetales;f__Micrococcaceae | 0 | 0 | 2 | 7 | 19 | 51 | 122.4 |
| k__Bacteria;p__AD3;c__JG37-AG-4;o__;f__ | 0 | 1 | 3 | 6 | 12 | 25.8 | 40.4 |
| k__Bacteria;p__WPS-2;c__;o__;f__ | 0 | 0 | 2 | 6 | 11 | 17 | 26 |
| k__Archaea;p__Crenarchaeota;c__MBGA;o__NRP-J;f__ | 0 | 1 | 3 | 6 | 11 | 19 | 40.4 |
| k__Bacteria;p__Acidobacteria;c__Solibacteres;o__Solibacterales;f__ | 0 | 1 | 2 | 5 | 9 | 13 | 20 |
| k__Bacteria;p__Nitrospirae;c__Nitrospira;o__Nitrospirales;f__Nitrospiraceae | 0 | 1 | 3 | 5 | 9 | 14.8 | 28 |
| k__Bacteria;p__Actinobacteria;c__Thermoleophilia;o__Solirubrobacterales;f__ | 0 | 1 | 2 | 5 | 8 | 12.8 | 17.4 |
| k__Bacteria;__;__;__;__ | 0 | 2 | 3 | 5 | 8 | 14 | 20.4 |
| k__Bacteria;p__Chloroflexi;c__Ktedonobacteria;o__Ktedonobacterales;f__Ktedonobacteraceae | 0 | 1 | 2 | 5 | 10 | 16 | 28.2 |
| k__Bacteria;p__Chloroflexi;c__Ktedonobacteria;o__TK10;f__ | 0 | 1 | 2 | 5 | 9 | 14 | 20 |
| k__Bacteria;p__Actinobacteria;c__Thermoleophilia;o__Solirubrobacterales;f__Conexibacteraceae | 0 | 1 | 3 | 5 | 7 | 10 | 12.8 |
| k__Bacteria;p__Nitrospirae;c__Nitrospira;o__Nitrospirales;f__0319-6A21 | 0 | 0 | 1 | 5 | 14 | 37 | 82.4 |
| k__Bacteria;p__Firmicutes;c__Bacilli;o__Bacillales;f__Paenibacillaceae | 0 | 0 | 2 | 4 | 7 | 12 | 42 |
| k__Bacteria;p__Chloroflexi;c__Ellin6529;o__;f__ | 0 | 0 | 1 | 4 | 10 | 16.8 | 38.2 |
| k__Bacteria;p__Proteobacteria;c__Alphaproteobacteria;o__Rhizobiales;f__Bradyrhizobiaceae | 0 | 0 | 2 | 4 | 11 | 21 | 28.4 |
| k__Bacteria;p__Proteobacteria;__;__;__ | 0 | 1 | 2 | 4 | 7 | 10.8 | 15.8 |
| k__Bacteria;p__Chloroflexi;__;__;__ | 0 | 0 | 2 | 4 | 9 | 13 | 20 |
| k__Bacteria;p__Actinobacteria;c__Acidimicrobiia;o__Acidimicrobiales;f__ | 0 | 1 | 3 | 4 | 6 | 9 | 11 |
| k__Bacteria;p__Planctomycetes;c__Phycisphaerae;o__WD2101;f__ | 0 | 0 | 1 | 3 | 5 | 8 | 13.8 |
| k__Bacteria;p__Actinobacteria;c__Acidimicrobiia;o__Acidimicrobiales;f__EB1017 | 0 | 0 | 1 | 3 | 5 | 8.8 | 14.2 |
| k__Bacteria;p__Proteobacteria;c__Betaproteobacteria;o__Burkholderiales;f__Burkholderiaceae | 0 | 0 | 1 | 3 | 7 | 23 | 85.4 |
| k__Bacteria;p__Proteobacteria;c__Gammaproteobacteria;o__Xanthomonadales;f__Xanthomonadaceae | 0 | 0 | 1 | 3 | 7 | 12 | 22.2 |
| k__Bacteria;p__Firmicutes;c__Bacilli;o__Bacillales;__ | 0 | 0 | 1 | 3 | 6 | 16 | 29 |

| | | | | | | | |
|---|---|---|---|---|---|---|---|
| k__Bacteria;p__Proteobacteria;c__Gammaproteobacteria;o__Xanthomonadales;f__Sinobacteraceae | 0 | 0 | 1 | 3 | 5 | 9.8 | 13 |
| k__Bacteria;p__Firmicutes;c__Bacilli;o__Bacillales;f__Bacillaceae | 0 | 0 | 1 | 3 | 6 | 15.8 | 26.8 |
| k__Bacteria;p__Verrucomicrobia;c__[Pedosphaerae];o__[Pedosphaerales];f__ | 0 | 0 | 0 | 2 | 5 | 9.8 | 21 |
| k__Bacteria;p__Actinobacteria;c__Actinobacteria;o__Actinomycetales;f__Thermomonosporaceae | 0 | 0 | 1 | 2 | 4 | 7 | 19.4 |
| k__Archaea;p__Crenarchaeota;c__Thaumarchaeota;o__Cenarchaeales;f__SAGMA-X | 0 | 0 | 0 | 2 | 5 | 12 | 26.4 |
| k__Bacteria;p__Proteobacteria;c__Alphaproteobacteria;o__Rhodospirillales;f__Acetobacteraceae | 0 | 0 | 1 | 2 | 4 | 6 | 10 |
| k__Bacteria;p__Acidobacteria;c__Acidobacteriia;o__Acidobacteriales;f__Acidobacteriaceae | 0 | 0 | 1 | 2 | 6 | 14 | 23.8 |
| k__Bacteria;p__Chloroflexi;c__TK10;o__B07_WMSP1;f__FFCH4570 | 0 | 0 | 1 | 2 | 3 | 5 | 8.4 |
| k__Bacteria;p__Planctomycetes;c__Planctomycetia;o__Pirellulales;f__Pirellulaceae | 0 | 0 | 1 | 2 | 4 | 7.8 | 12.8 |
| k__Bacteria;p__Chloroflexi;c__Ktedonobacteria;o__JG30-KF-AS9;f__ | 0 | 0 | 1 | 2 | 5 | 10 | 21 |
| k__Bacteria;p__Proteobacteria;c__Alphaproteobacteria;o__Ellin329;f__ | 0 | 0 | 1 | 2 | 3 | 5 | 9 |
| k__Bacteria;p__Proteobacteria;c__Alphaproteobacteria;o__Sphingomonadales;f__Sphingomonadaceae | 0 | 0 | 1 | 2 | 7 | 20 | 39 |
| k__Bacteria;p__Proteobacteria;c__Deltaproteobacteria;o__Myxococcales;f__ | 0 | 0 | 0 | 2 | 4 | 7.8 | 11.4 |
| k__Bacteria;p__Proteobacteria;c__Betaproteobacteria;o__SC-I-84;f__ | 0 | 0 | 1 | 2 | 6 | 13.8 | 20.4 |
| k__Bacteria;p__Chloroflexi;c__Anaerolineae;o__;f__ | 0 | 0 | 0 | 2 | 4 | 8 | 15.4 |
| k__Bacteria;p__Verrucomicrobia;c__[Pedosphaerae];o__[Pedosphaerales];f__Ellin515 | 0 | 0 | 1 | 2 | 5 | 7 | 10 |
| k__Bacteria;p__Proteobacteria;c__Betaproteobacteria;o__Burkholderiales;f__Oxalobacteraceae | 0 | 0 | 1 | 2 | 4 | 10.8 | 20 |
| k__Bacteria;p__Planctomycetes;c__Planctomycetia;o__Planctomycetales;f__Planctomycetaceae | 0 | 0 | 0 | 2 | 3 | 4 | 6 |
| k__Bacteria;p__Bacteroidetes;c__[Saprospirae];o__[Saprospirales];f__Chitinophagaceae | 0 | 0 | 0 | 2 | 6 | 15.8 | 30.8 |
| k__Bacteria;p__Actinobacteria;c__Actinobacteria;o__Actinomycetales;f__Intrasporangiaceae | 0 | 0 | 0 | 1 | 4 | 10 | 16.8 |
| k__Bacteria;p__Proteobacteria;c__Betaproteobacteria;o__Ellin6067;f__ | 0 | 0 | 0 | 1 | 2 | 4 | 7 |
| k__Bacteria;p__Actinobacteria;c__Actinobacteria;o__Actinomycetales;f__Mycobacteriaceae | 0 | 0 | 0 | 1 | 2 | 5 | 7 |
| k__Bacteria;p__Chloroflexi;c__Ktedonobacteria;o__;f__ | 0 | 0 | 0 | 1 | 3 | 7 | 12 |
| k__Bacteria;p__Chloroflexi;c__P2-11E;o__;f__ | 0 | 0 | 0 | 1 | 2 | 4 | 7 |
| k__Bacteria;p__Actinobacteria;c__Actinobacteria;o__Actinomycetales;f__Nocardioidaceae | 0 | 0 | 0 | 1 | 5 | 10 | 18.4 |
| k__Bacteria;p__Actinobacteria;c__MB-A2-108;o__;f__ | 0 | 0 | 0 | 1 | 2 | 4 | 7 |
| k__Bacteria;p__Proteobacteria;c__Betaproteobacteria;o__Burkholderiales;f__Comamonadaceae | 0 | 0 | 0 | 1 | 5 | 11 | 22.4 |
| k__Bacteria;p__Proteobacteria;c__Alphaproteobacteria;o__Rhizobiales;f__ | 0 | 0 | 0 | 1 | 2 | 5 | 7 |
| k__Bacteria;p__Chloroflexi;c__Ktedonobacteria;o__Elev-1554;f__ | 0 | 0 | 0 | 1 | 2 | 4 | 7 |
| k__Bacteria;p__Firmicutes;c__Bacilli;o__Bacillales;f__Planococcaceae | 0 | 0 | 0 | 1 | 3 | 7 | 12 |
| k__Bacteria;p__Gemmatimonadetes;c__Gemmatimonadetes;o__N1423WL;f__ | 0 | 0 | 0 | 1 | 2 | 5 | 8 |
| k__Bacteria;p__Gemmatimonadetes;c__Gemmatimonadetes;o__Ellin5290;f__ | 0 | 0 | 0 | 1 | 3 | 5 | 8 |
| k__Bacteria;p__Chloroflexi;c__S085;o__;f__ | 0 | 0 | 0 | 1 | 3 | 8 | 11.4 |
| k__Bacteria;p__Proteobacteria;c__Alphaproteobacteria;o__Caulobacterales;f__Caulobacteraceae | 0 | 0 | 0 | 1 | 2 | 3 | 6 |
| k__Bacteria;p__Gemmatimonadetes;c__Gemmatimonadetes;o__;f__ | 0 | 0 | 0 | 1 | 3 | 7 | 13 |
| k__Bacteria;p__Actinobacteria;c__Actinobacteria;o__Actinomycetales;f__Micromonosporaceae | 0 | 0 | 0 | 1 | 2 | 7 | 15.4 |
| k__Bacteria;p__Chloroflexi;c__TK17;o__;f__ | 0 | 0 | 0 | 1 | 2 | 4 | 5 |
| k__Bacteria;p__Acidobacteria;c__iii1-8;o__DS-18;f__ | 0 | 0 | 0 | 1 | 4 | 8 | 12 |
| k__Bacteria;p__Proteobacteria;c__Gammaproteobacteria;o__Legionellales;f__Coxiellaceae | 0 | 0 | 0 | 1 | 1 | 3 | 5 |

| | | | | | | | |
|---|---|---|---|---|---|---|---|
| k__Bacteria;p__Proteobacteria;c__Alphaproteobacteria;o__Rhodospirillales;f__Rhodospirillaceae | 7 | 12 | 19 | 27 | 39 | 49.96 | 64.88 |

| Feature ID | 2% | 9% | 25% | 50% | 75% | 91% | 98% |
|---|---|---|---|---|---|---|---|
| k__Bacteria;p__A | 13.5 | 30 | 47 | 68 | 90 | 109.25 | 126 |
| k__Bacteria;p__F | 6 | 11 | 17 | 27 | 39 | 49 | 54 |

| Feature ID | 2% | 9% | 25% | 50% | 75% | 91% | 98% |
|---|---|---|---|---|---|---|---|
| k__Bacteria;p__( | 54.6 | 94.4 | 148 | 223 | 309 | 414.4 | 592.6 |
| k__Bacteria;p__/ | 17.6 | 29.2 | 70 | 117 | 203 | 287 | 400 |
| k__Bacteria;p__/ | 27.6 | 40 | 55 | 77 | 113 | 145 | 174.4 |
| k__Bacteria;p__/ | 7 | 13 | 24 | 48 | 82 | 122.8 | 162.8 |
| k__Bacteria;p__/ | 5.6 | 13 | 23 | 36 | 53 | 71.8 | 108 |
| k__Bacteria;p__\ | 4.6 | 12 | 24 | 36 | 56 | 88 | 125 |
| k__Bacteria;p__/ | 11 | 17.2 | 24 | 33 | 46 | 63 | 78.8 |
| k__Bacteria;p__I | 8.6 | 14 | 20 | 28 | 37 | 53.8 | 75.4 |
| k__Bacteria;p__I | 6 | 10.2 | 16 | 22 | 30 | 39 | 54.4 |
| k__Bacteria;p__I | 6.6 | 9 | 13 | 20 | 29 | 41 | 57 |
