## Supplemental Figures for "Soil depth determines the microbial communities in *Sorghum bicolor* fields"

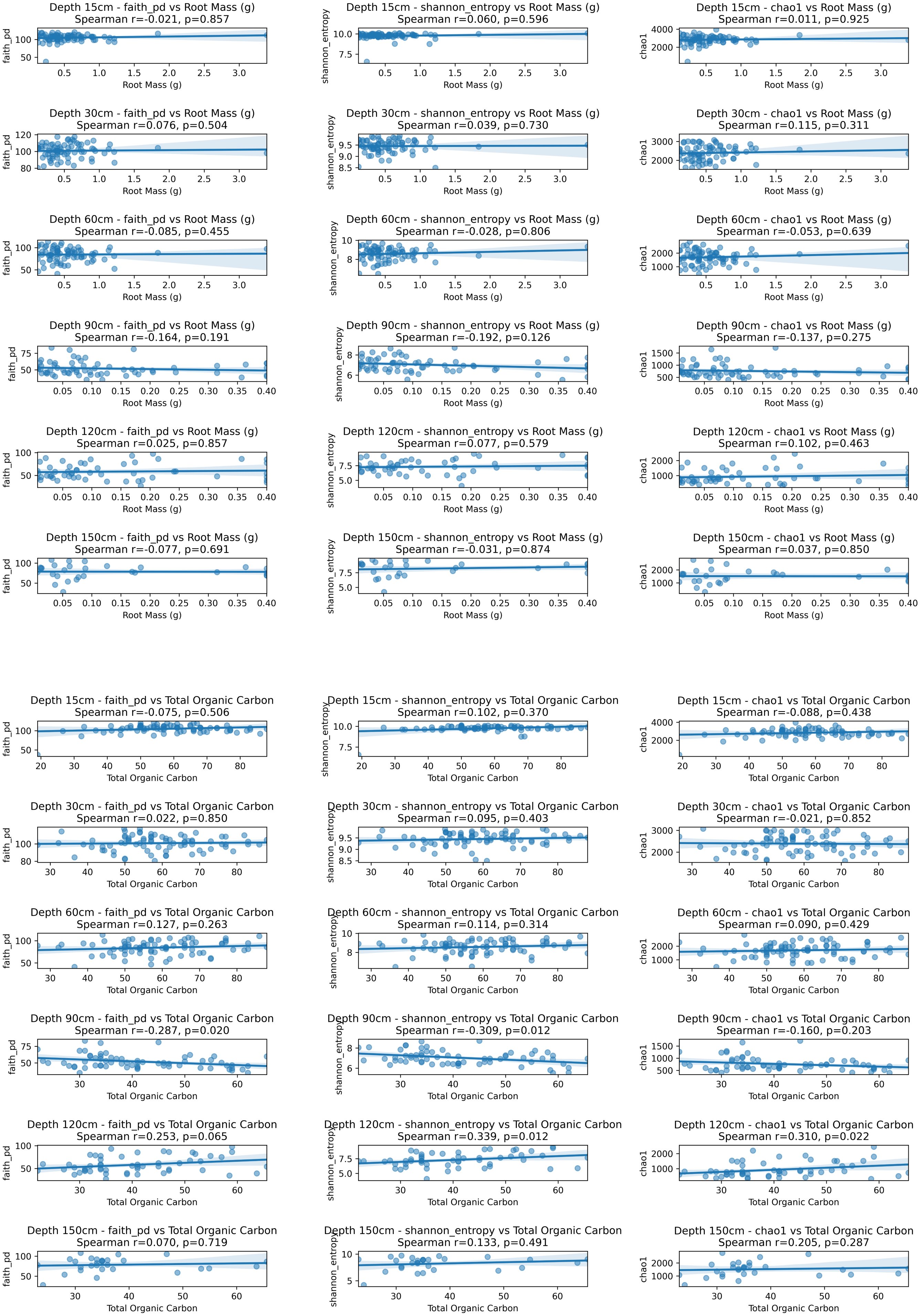


Fig S1: Changes in alpha diversity levels with root mass and total organic carbon at each depth displaying Spearman's rank correlation coefficients and p-values in the plot titles.

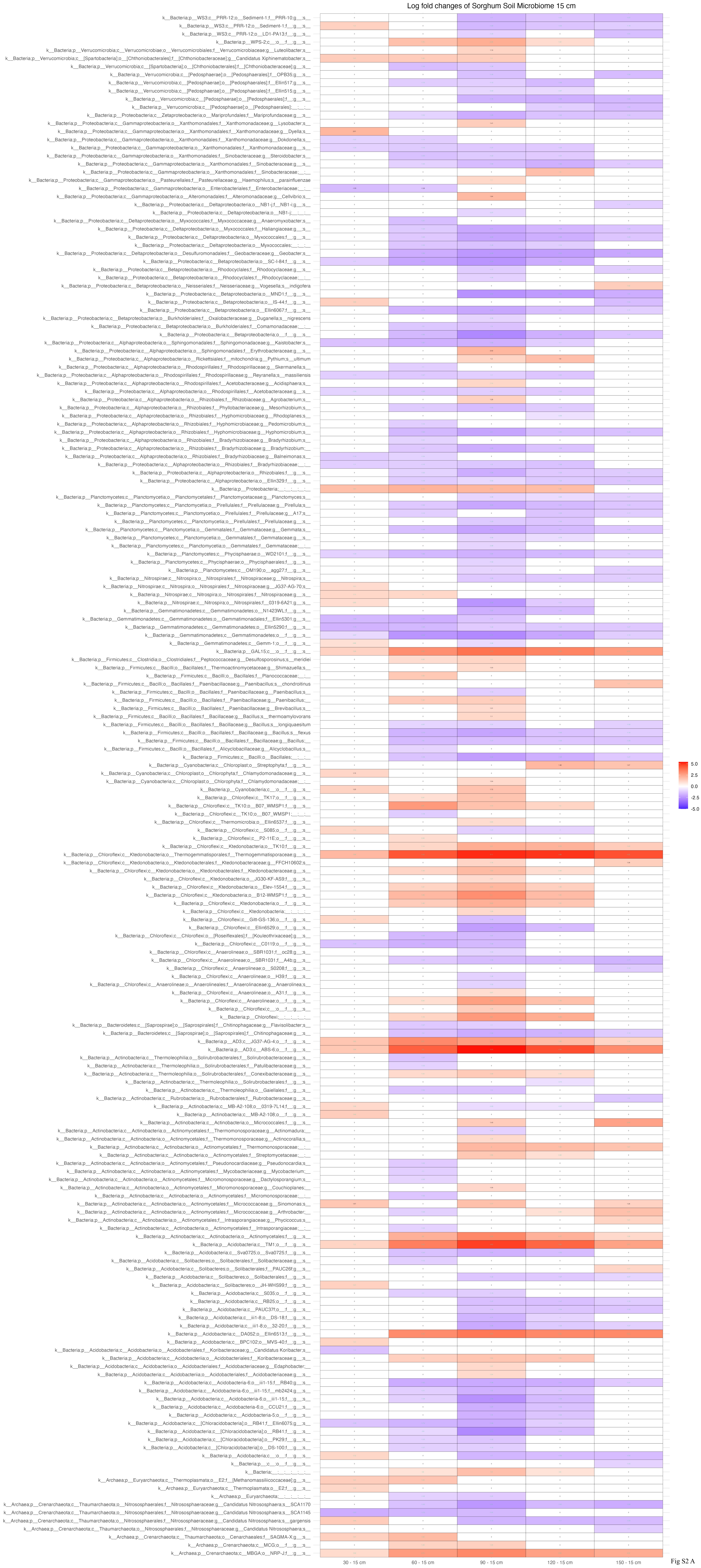
Log fold changes of Sorghum Soil Microbiome 15 cm
30 - 15 cm
60 - 15 cm
90 - 15 cm
120 - 15 cm
150 - 15 cm
5.0
2.5
0.0
-2.5
-5.0

Fig S2 A

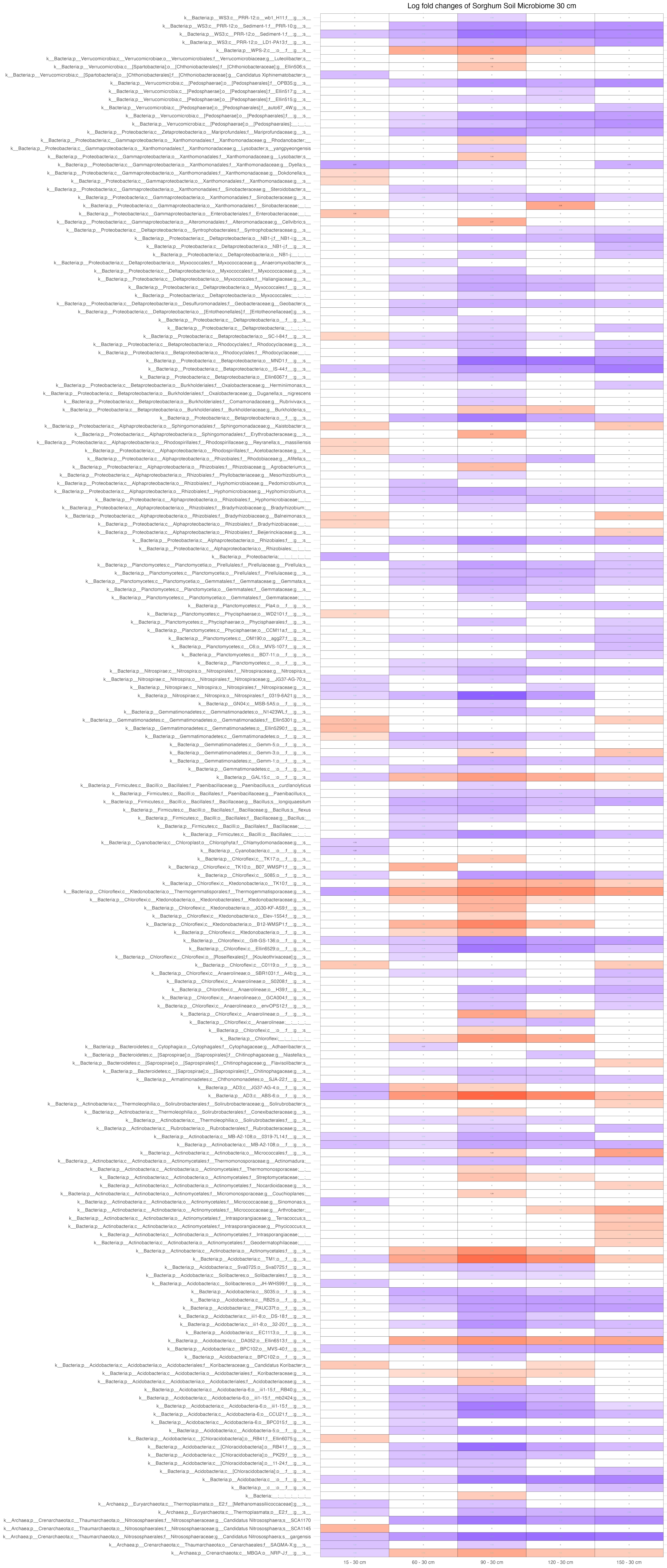
Log fold changes of Sorghum Soil Microbiome 30 cm
15 - 30 cm
60 - 30 cm
90 - 30 cm
120 - 30 cm
150 - 30 cm
5.0
2.5
0.0
-2.5
-5.0

Fig S2 B

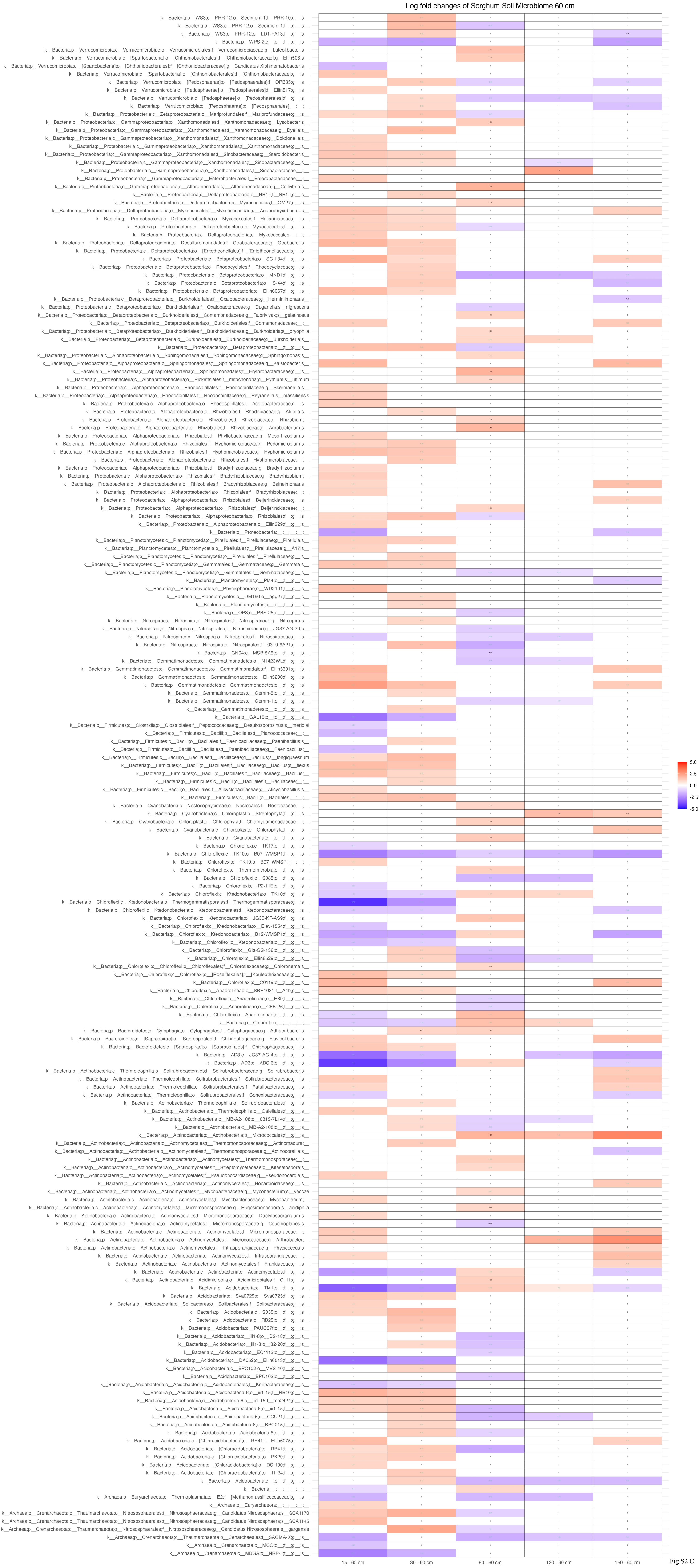
Log fold changes of Sorghum Soil Microbiome 60 cm
15 - 60 cm
30 - 60 cm
90 - 60 cm
120 - 60 cm
150 - 60 cm
5.0
2.5
0.0
-2.5
-5.0
Fig S2 C

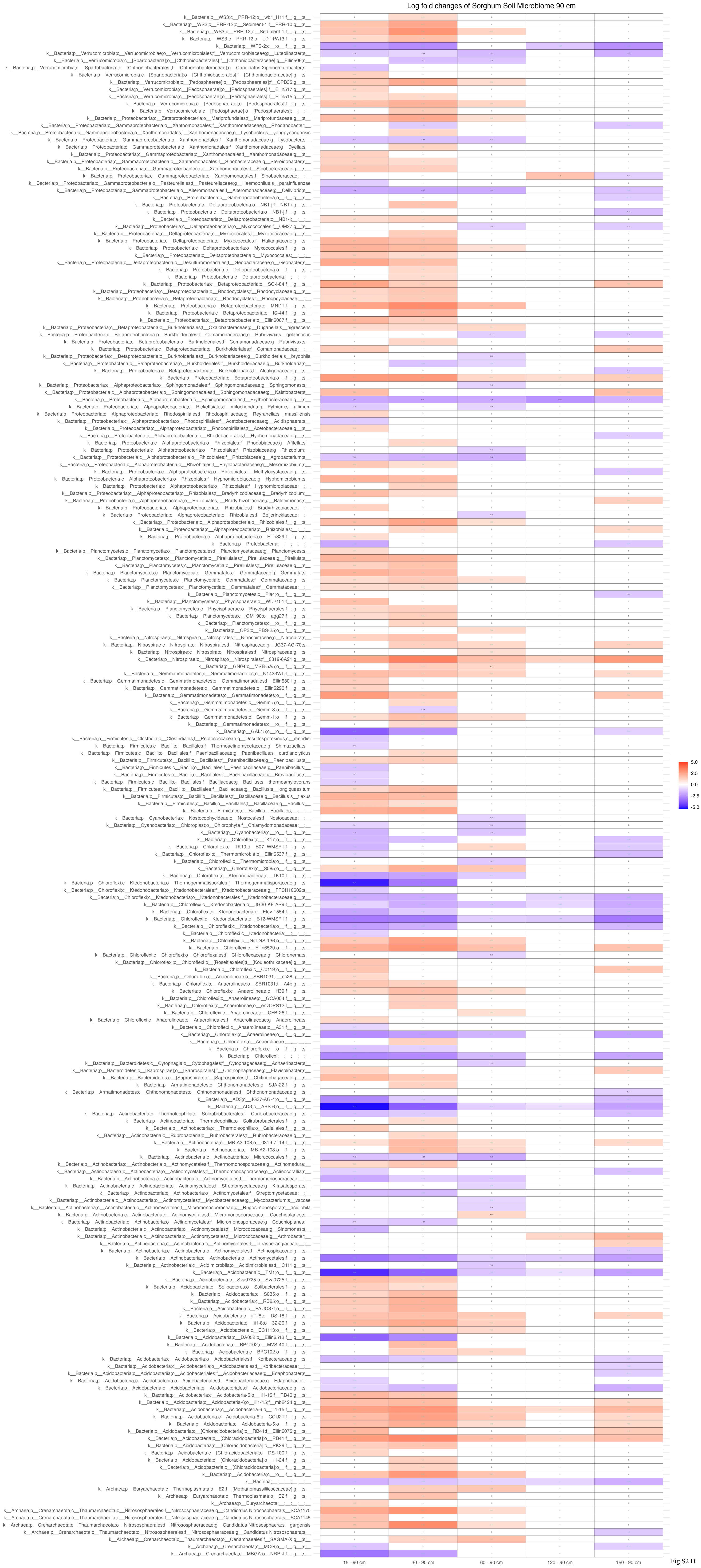
Log fold changes of Sorghum Soil Microbiome 90 cm
15 - 90 cm
30 - 90 cm
60 - 90 cm
120 - 90 cm
150 - 90 cm
5.0
2.5
0.0
-2.5
-5.0

Fig S2 D

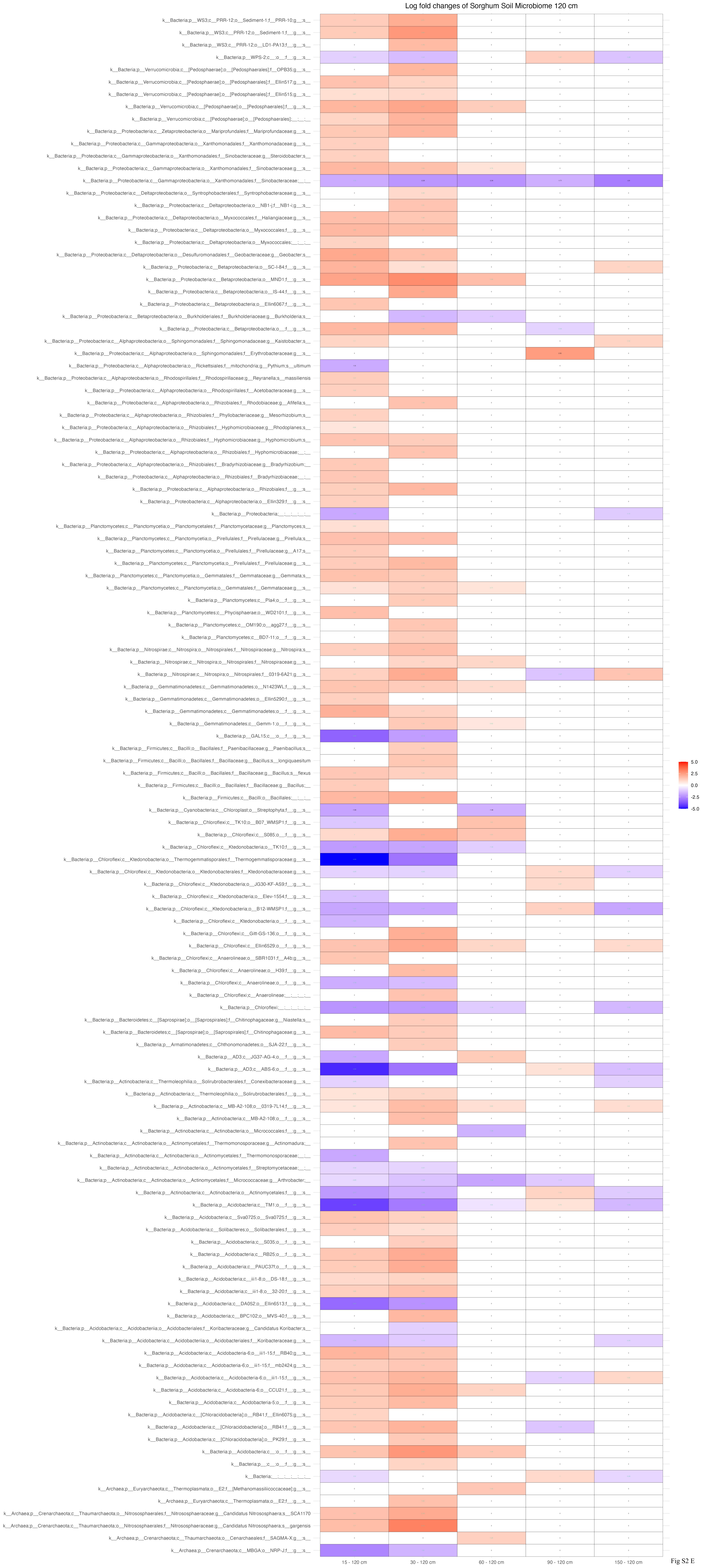
Log fold changes of Sorghum Soil Microbiome 120 cm
k__Bacteria;p__WS3;c__PRR-12;o__Sediment-1;f__PRR-10;g__;s__
k__Bacteria;p__WS3;c__PRR-12;o__Sediment-1;f__;g__;s__
k__Bacteria;p__WS3;c__PRR-12;o__LD1-PA13;f__;g__;s__
k__Bacteria;p__WPS-2;c__;o__;f__;g__;s__
k__Bacteria;p__Verrucomicrobia;c__[Pedosphaerae];o__[Pedosphaerales];f__OPB35;g__;s__
k__Bacteria;p__Verrucomicrobia;c__[Pedosphaerae];o__[Pedosphaerales];f__Ellin517;g__;s__
k__Bacteria;p__Verrucomicrobia;c__[Pedosphaerae];o__[Pedosphaerales];f__Ellin515;g__;s__
k__Bacteria;p__Verrucomicrobia;c__[Pedosphaerae];o__[Pedosphaerales];f__;g__;s__
k__Bacteria;p__Verrucomicrobia;c__[Pedosphaerae];o__[Pedosphaerales];__;__;__
k__Bacteria;p__Proteobacteria;c__Zetaproteobacteria;o__Mariprofundales;f__Mariprofundaceae;g__;s__
k__Bacteria;p__Proteobacteria;c__Gammaproteobacteria;o__Xanthomonadales;f__Xanthomonadaceae;g__;s__
k__Bacteria;p__Proteobacteria;c__Gammaproteobacteria;o__Xanthomonadales;f__Sinobacteraceae;g__Steroidobacter;s__
k__Bacteria;p__Proteobacteria;c__Gammaproteobacteria;o__Xanthomonadales;f__Sinobacteraceae;g__;s__
k__Bacteria;p__Proteobacteria;c__Gammaproteobacteria;o__Xanthomonadales;f__Sinobacteraceae;__;__
k__Bacteria;p__Proteobacteria;c__Deltaproteobacteria;o__Syntrophobacterales;f__Syntrophobacteraceae;g__;s__
k__Bacteria;p__Proteobacteria;c__Deltaproteobacteria;o__NB1-j;f__NB1-i;g__;s__
k__Bacteria;p__Proteobacteria;c__Deltaproteobacteria;o__Myxococcales;f__Haliangiaceae;g__;s__
k__Bacteria;p__Proteobacteria;c__Deltaproteobacteria;o__Myxococcales;f__;g__;s__
k__Bacteria;p__Proteobacteria;c__Deltaproteobacteria;o__Myxococcales;__;__;__
k__Bacteria;p__Proteobacteria;c__Deltaproteobacteria;o__Desulfuromonadales;f__Geobacteraceae;g__Geobacter;s__
k__Bacteria;p__Proteobacteria;c__Betaproteobacteria;o__SC-I-84;f__;g__;s__
k__Bacteria;p__Proteobacteria;c__Betaproteobacteria;o__MND1;f__;g__;s__
k__Bacteria;p__Proteobacteria;c__Betaproteobacteria;o__IS-44;f__;g__;s__
k__Bacteria;p__Proteobacteria;c__Betaproteobacteria;o__Ellin6067;f__;g__;s__
k__Bacteria;p__Proteobacteria;c__Betaproteobacteria;o__Burkholderiales;f__Burkholderiaceae;g__Burkholderia;s__
k__Bacteria;p__Proteobacteria;c__Betaproteobacteria;o__;f__;g__;s__
k__Bacteria;p__Proteobacteria;c__Alphaproteobacteria;o__Sphingomonadales;f__Sphingomonadaceae;g__Kaistobacter;s__
k__Bacteria;p__Proteobacteria;c__Alphaproteobacteria;o__Sphingomonadales;f__Erythrobacteraceae;g__;s__
k__Bacteria;p__Proteobacteria;c__Alphaproteobacteria;o__Rickettsiales;f__mitochondria;g__Pythium;s__ultimum
k__Bacteria;p__Proteobacteria;c__Alphaproteobacteria;o__Rhodospirillales;f__Rhodospirillaceae;g__Reyranella;s__massiliensis
k__Bacteria;p__Proteobacteria;c__Alphaproteobacteria;o__Rhodospirillales;f__Acetobacteraceae;g__;s__
k__Bacteria;p__Proteobacteria;c__Alphaproteobacteria;o__Rhizobiales;f__Rhodobiaceae;g__Afifella;s__
k__Bacteria;p__Proteobacteria;c__Alphaproteobacteria;o__Rhizobiales;f__Phyllobacteriaceae;g__Mesorhizobium;s__
k__Bacteria;p__Proteobacteria;c__Alphaproteobacteria;o__Rhizobiales;f__Hyphomicrobiaceae;g__Rhodoplanes;s__
k__Bacteria;p__Proteobacteria;c__Alphaproteobacteria;o__Rhizobiales;f__Hyphomicrobiaceae;g__Hyphomicrobium;s__
k__Bacteria;p__Proteobacteria;c__Alphaproteobacteria;o__Rhizobiales;f__Hyphomicrobiaceae;__;__
k__Bacteria;p__Proteobacteria;c__Alphaproteobacteria;o__Rhizobiales;f__Bradyrhizobiaceae;g__Bradyrhizobium;__
k__Bacteria;p__Proteobacteria;c__Alphaproteobacteria;o__Rhizobiales;f__Bradyrhizobiaceae;__;__
k__Bacteria;p__Proteobacteria;c__Alphaproteobacteria;o__Rhizobiales;f__;g__;s__
k__Bacteria;p__Proteobacteria;c__Alphaproteobacteria;o__Ellin329;f__;g__;s__
k__Bacteria;p__Proteobacteria;__;__;__;__;__
k__Bacteria;p__Planctomycetes;c__Planctomycetia;o__Planctomycetales;f__Planctomycetaceae;g__Planctomyces;s__
k__Bacteria;p__Planctomycetes;c__Planctomycetia;o__Pirellulales;f__Pirellulaceae;g__Pirellula;s__
k__Bacteria;p__Planctomycetes;c__Planctomycetia;o__Pirellulales;f__Pirellulaceae;g__A17;s__
k__Bacteria;p__Planctomycetes;c__Planctomycetia;o__Pirellulales;f__Pirellulaceae;g__;s__
k__Bacteria;p__Planctomycetes;c__Planctomycetia;o__Gemmatales;f__Gemmataceae;g__Gemmata;s__
k__Bacteria;p__Planctomycetes;c__Planctomycetia;o__Gemmatales;f__Gemmataceae;g__;s__
k__Bacteria;p__Planctomycetes;c__Pla4;o__;f__;g__;s__
k__Bacteria;p__Planctomycetes;c__Phycisphaerae;o__WD2101;f__;g__;s__
k__Bacteria;p__Planctomycetes;c__OM190;o__agg27;f__;g__;s__
k__Bacteria;p__Planctomycetes;c__BD7-11;o__;f__;g__;s__
k__Bacteria;p__Nitrospirae;c__Nitrospira;o__Nitrospirales;f__Nitrospiraceae;g__Nitrospira;s__
k__Bacteria;p__Nitrospirae;c__Nitrospira;o__Nitrospirales;f__Nitrospiraceae;g__;s__
k__Bacteria;p__Nitrospirae;c__Nitrospira;o__Nitrospirales;f__0319-6A21;g__;s__
k__Bacteria;p__Gemmatimonadetes;c__Gemmatimonadetes;o__N1423WL;f__;g__;s__
k__Bacteria;p__Gemmatimonadetes;c__Gemmatimonadetes;o__Ellin5290;f__;g__;s__
k__Bacteria;p__Gemmatimonadetes;c__Gemmatimonadetes;o__;f__;g__;s__
k__Bacteria;p__Gemmatimonadetes;c__Gemm-1;o__;f__;g__;s__
k__Bacteria;p__GAL15;c__;o__;f__;g__;s__
k__Bacteria;p__Firmicutes;c__Bacilli;o__Bacillales;f__Paenibacillaceae;g__Paenibacillus;s__
k__Bacteria;p__Firmicutes;c__Bacilli;o__Bacillales;f__Bacillaceae;g__Bacillus;s__longiquaesitum
k__Bacteria;p__Firmicutes;c__Bacilli;o__Bacillales;f__Bacillaceae;g__Bacillus;s__flexus
k__Bacteria;p__Firmicutes;c__Bacilli;o__Bacillales;f__Bacillaceae;g__Bacillus;__
k__Bacteria;p__Firmicutes;c__Bacilli;o__Bacillales;__;__;__
k__Bacteria;p__Cyanobacteria;c__Chloroplast;o__Streptophyta;f__;g__;s__
k__Bacteria;p__Chloroflexi;c__TK10;o__B07_WMSP1;f__;g__;s__
k__Bacteria;p__Chloroflexi;c__S085;o__;f__;g__;s__
k__Bacteria;p__Chloroflexi;c__Ktedonobacteria;o__TK10;f__;g__;s__
k__Bacteria;p__Chloroflexi;c__Ktedonobacteria;o__Thermogemmatisporales;f__Thermogemmatisporaceae;g__;s__
k__Bacteria;p__Chloroflexi;c__Ktedonobacteria;o__Ktedonobacterales;f__Ktedonobacteraceae;g__;s__
k__Bacteria;p__Chloroflexi;c__Ktedonobacteria;o__JG30-KF-AS9;f__;g__;s__
k__Bacteria;p__Chloroflexi;c__Ktedonobacteria;o__Elev-1554;f__;g__;s__
k__Bacteria;p__Chloroflexi;c__Ktedonobacteria;o__B12-WMSP1;f__;g__;s__
k__Bacteria;p__Chloroflexi;c__Ktedonobacteria;o__;f__;g__;s__
k__Bacteria;p__Chloroflexi;c__Gitt-GS-136;o__;f__;g__;s__
k__Bacteria;p__Chloroflexi;c__Ellin6529;o__;f__;g__;s__
k__Bacteria;p__Chloroflexi;c__Anaerolineae;o__SBR1031;f__A4b;g__;s__
k__Bacteria;p__Chloroflexi;c__Anaerolineae;o__H39;f__;g__;s__
k__Bacteria;p__Chloroflexi;c__Anaerolineae;o__;f__;g__;s__
k__Bacteria;p__Chloroflexi;c__Anaerolineae;__;__;__;__
k__Bacteria;p__Chloroflexi;__;__;__;__;__
k__Bacteria;p__Bacteroidetes;c__[Saprospirae];o__[Saprospirales];f__Chitinophagaceae;g__Niastella;s__
k__Bacteria;p__Bacteroidetes;c__[Saprospirae];o__[Saprospirales];f__Chitinophagaceae;g__;s__
k__Bacteria;p__Armatimonadetes;c__Chthonomonadetes;o__SJA-22;f__;g__;s__
k__Bacteria;p__AD3;c__JG37-AG-4;o__;f__;g__;s__
k__Bacteria;p__AD3;c__ABS-6;o__;f__;g__;s__
k__Bacteria;p__Actinobacteria;c__Thermoleophilia;o__Solirubrobacterales;f__Conexibacteraceae;g__;s__
k__Bacteria;p__Actinobacteria;c__Thermoleophilia;o__Solirubrobacterales;f__;g__;s__
k__Bacteria;p__Actinobacteria;c__MB-A2-108;o__0319-7L14;f__;g__;s__
k__Bacteria;p__Actinobacteria;c__MB-A2-108;o__;f__;g__;s__
k__Bacteria;p__Actinobacteria;c__Actinobacteria;o__Micrococcales;f__;g__;s__
k__Bacteria;p__Actinobacteria;c__Actinobacteria;o__Actinomycetales;f__Thermomonosporaceae;g__Actinomadura;__
k__Bacteria;p__Actinobacteria;c__Actinobacteria;o__Actinomycetales;f__Thermomonosporaceae;__;__
k__Bacteria;p__Actinobacteria;c__Actinobacteria;o__Actinomycetales;f__Streptomycetaceae;__;__
k__Bacteria;p__Actinobacteria;c__Actinobacteria;o__Actinomycetales;f__Micrococcaceae;g__Arthrobacter;__
k__Bacteria;p__Actinobacteria;c__Actinobacteria;o__Actinomycetales;f__;g__;s__
k__Bacteria;p__Acidobacteria;c__TM1;o__;f__;g__;s__
k__Bacteria;p__Acidobacteria;c__Sva0725;o__Sva0725;f__;g__;s__
k__Bacteria;p__Acidobacteria;c__Solibacteres;o__Solibacterales;f__;g__;s__
k__Bacteria;p__Acidobacteria;c__S035;o__;f__;g__;s__
k__Bacteria;p__Acidobacteria;c__RB25;o__;f__;g__;s__
k__Bacteria;p__Acidobacteria;c__PAUC37f;o__;f__;g__;s__
k__Bacteria;p__Acidobacteria;c__iii1-8;o__DS-18;f__;g__;s__
k__Bacteria;p__Acidobacteria;c__iii1-8;o__32-20;f__;g__;s__
k__Bacteria;p__Acidobacteria;c__DA052;o__Ellin6513;f__;g__;s__
k__Bacteria;p__Acidobacteria;c__BPC102;o__MVS-40;f__;g__;s__
k__Bacteria;p__Acidobacteria;c__Acidobacteria;o__Acidobacteriales;f__Koribacteraceae;g__Candidatus Koribacter;s__
k__Bacteria;p__Acidobacteria;c__Acidobacteriia;o__Acidobacteriales;f__Koribacteraceae;g__;s__
k__Bacteria;p__Acidobacteria;c__Acidobacteria-6;o__iii1-15;f__RB40;g__;s__
k__Bacteria;p__Acidobacteria;c__Acidobacteria-6;o__iii1-15;f__mb2424;g__;s__
k__Bacteria;p__Acidobacteria;c__Acidobacteria-6;o__iii1-15;f__;g__;s__
k__Bacteria;p__Acidobacteria;c__Acidobacteria-6;o__CCU21;f__;g__;s__
k__Bacteria;p__Acidobacteria;c__Acidobacteria-5;o__;f__;g__;s__
k__Bacteria;p__Acidobacteria;c__[Chloracidobacteria];o__RB41;f__Ellin6075;g__;s__
k__Bacteria;p__Acidobacteria;c__[Chloracidobacteria];o__RB41;f__;g__;s__
k__Bacteria;p__Acidobacteria;c__[Chloracidobacteria];o__PK29;f__;g__;s__
k__Bacteria;p__Acidobacteria;c__;o__;f__;g__;s__
k__Bacteria;p__;c__;o__;f__;g__;s__
k__Bacteria;__;__;__;__;__;__
k__Archaea;p__Euryarchaeota;c__Thermoplasmata;o__E2;f__[Methanomassiliicoccaceae];g__;s__
k__Archaea;p__Euryarchaeota;c__Thermoplasmata;o__E2;f__;g__;s__
k__Archaea;p__Crenarchaeota;c__Thaumarchaeota;o__Nitrososphaerales;f__Nitrososphaeraceae;g__Candidatus Nitrososphaera;s__SCA1170
k__Archaea;p__Crenarchaeota;c__Thaumarchaeota;o__Nitrososphaerales;f__Nitrososphaeraceae;g__Candidatus Nitrososphaera;s__gargensis
k__Archaea;p__Crenarchaeota;c__Thaumarchaeota;o__Cenarchaeales;f__SAGMA-X;g__;s__
k__Archaea;p__Crenarchaeota;c__MBGA;o__NRP-J;f__;g__;s__
15 - 120 cm
30 - 120 cm
60 - 120 cm
90 - 120 cm
150 - 120 cm
5.0
2.5
0.0
-2.5
-5.0

Fig S2 E

Log fold changes of Sorghum Soil Microbiome 150 cm

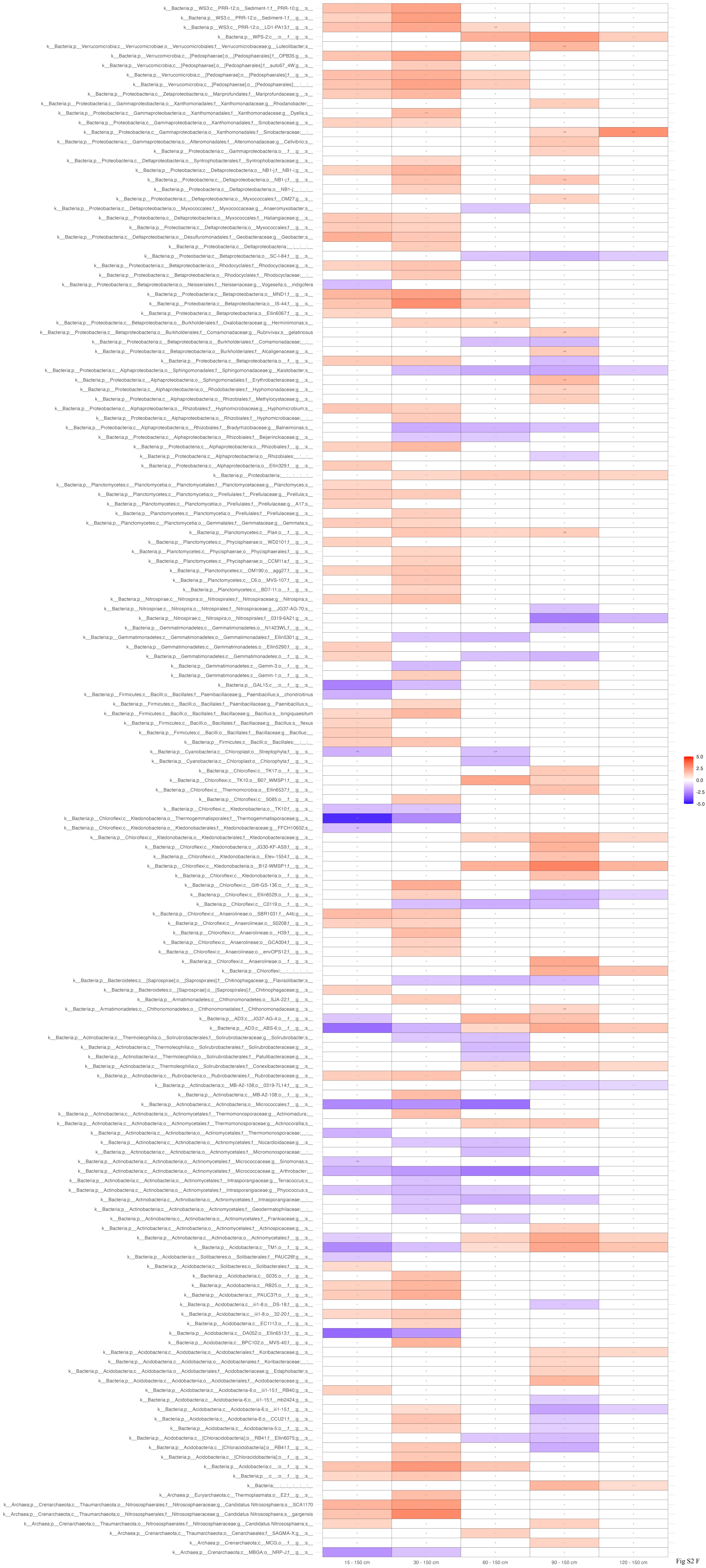


Fig S2 F

**Fig S2:** Log fold changes of microbial species at various depths. A-F***)*** Heatmaps displaying all of the log fold changes (LFC) of the *S. bicolor* soil microbiome across at different depths compared to the reference depth A) 15 cm, B) 30 cm, C) 60 cm, D) 90 cm, E) 120 cm and F) 150 cm. The y-axes display the taxa arranged in alphabetical order. The x-axes show the different soil depth comparisons. The color gradient ranges from blue, indicating negative log fold changes, to red, indicating positive log fold changes, with white representing no change. The midpoint of the scale is set to 0, with limits ranging from -5.4 to 5.4. Comparisons that have successfully passed the sensitivity analysis for pseudo-count addition are denoted by aquamarine text. Taxa with no significant values in any comparisons were excluded to focus on the relevant data.
